# Mitochondrial Signaling: Nitric Oxide Synthesis by Cytochrome c Oxidase and Its Oxygen Sensitivity Are Modulated by Adenine Nucleotides

**DOI:** 10.64898/2026.08.09.743791

**Authors:** Pablo R. Castello, Kerri A. Ball, Robert O. Poyton

## Abstract

Nitrite can be reduced to nitric oxide (NO) by several heme- and molybdenum-containing proteins, including mitochondrial cytochrome c oxidase (Cco). This activity, designated Cco/NO, has been implicated in hypoxic signaling, but its regulation and quantitative significance relative to other NO-producing systems remain uncertain. We examined its modulation by adenine nucleotides using detergent-solubilized yeast and mouse brain mitochondria supplied with 1 mM nitrite and an ascorbate/TMPD/cytochrome c electron-donor system. ADP and ATP differentially modulated Cco/NO activity, and ADP shifted the apparent oxygen sensitivity of Cco/NO toward lower O₂, with significant stimulation at 10 µM O₂ and below. Nucleotide regulation was also isoform-dependent: ATP slightly inhibited Va-containing Cco but strongly stimulated Vb-containing Cco under anoxic conditions. Rates normalized to cytochrome aa₃ demonstrate multi-turnover nitrite-reductase capacity under these substrate-driven assay conditions. Both the cellular ADP/ATP ratio and subsequently assayed Cco/NO activity increased transiently following a hypoxic shift. These findings establish metabolic and isoform-dependent gating of the catalytic capacity of Cco/NO; they do not establish its fractional contribution to total cellular NO or its operation at physiological nitrite concentrations in intact, coupled mitochondria.

## 1. INTRODUCTION

Nitric oxide (NO) is a short-lived signaling molecule whose biological actions depend on the local balance among NO production, consumption, target sensitivity, and compartmental diffusion [1]. NO and reactive nitrogen species participate in diverse physiological and stress-response pathways [2–7]. Although nitric oxide synthases catalyze the canonical oxygen-dependent conversion of L-arginine to NO, nitrate and nitrite provide a parallel redox reservoir for NO bioactivity. The nitrate–nitrite–NO axis is especially relevant during hypoxia and acidosis, when heme proteins, molybdenum enzymes, mitochondrial redox centers, and proton-coupled chemistry can contribute to nitrite reduction and NO bioavailability [8–13].

Cytochrome c oxidase (Cco), the terminal enzyme of the mitochondrial electron-transport chain, is one such nitrite-reducing system. Previous work from this group showed that Cco can generate NO from nitrite under low-oxygen conditions, a reaction designated Cco/NO, and implicated this activity in hypoxic signaling in yeast and mammalian cells [14, 15]. The oxygen-regulated subunit V isoforms of yeast Cco (Va and Vb) differ in the oxygen tensions that permit NO formation, with the hypoxia-associated Vb isoform supporting NO synthesis at higher O₂ concentrations than the aerobic Va isoform [16]. Low-intensity light can further stimulate nitrite-dependent NO synthesis by Cco without increasing O₂ consumption [17, 18]. Nitrite reduction is not the only reaction the binuclear heme a₃–Cu_B_ center supports besides the four-electron reduction of O₂ to water: the same site binds and turns over other small gaseous and anionic ligands, and its interconversion with NO in particular is well characterized [19–23]. The enzyme also oxygenates carbon monoxide to carbon dioxide, an activity demonstrated both with the purified bovine enzyme and in intact mitochondria [24, 25]. Cco/NO should therefore be read as one member of a set of non-canonical reactions at that center, not as evidence that the enzyme is a dedicated two-state device. Together, these observations establish Cco/NO as a regulated biochemical output rather than a fixed stoichiometric by-product of respiratory inhibition.

The conversion of NO₂⁻ to NO most likely takes place on the binuclear reaction center (heme a₃–Cu_B_) of Cco. Evidence for this comes from the findings: that a NO₂⁻– ferric a₃ complex [19–22] and a heme-nitrosyl complex a₃Z⁺–NO [23] are formed at the binuclear center of Cco in the presence of NO₂⁻; that 590 nm light, which overlaps the absorption spectrum of the heme a₃Z⁺–NO complex, stimulates Cco/NO activity [18]; and, more recently, that an inorganic reduced heme–copper assembly, resembling the Cco binuclear reaction center, reduces NO₂⁻ to NO [26]. In that model system the ferrous heme is the reductant, but the cupric ion is required: nitrite bound to a ferric heme did not react when a cuprous complex was added, and the fully reduced metal pair showed no nitrite-reductase activity. Two equivalents of ferrous heme react with one equivalent of the copper(II)-nitrito complex to give a 1:1 mixture of the heme-nitrosyl and µ-oxo products, the second equivalent being consumed by the high affinity of NO for ferrous heme. The same assembly also performs the reverse reaction, oxidizing NO back to nitrite in 95% yield [26]. That work was done on synthetic complexes in acetone rather than in water or on an enzyme, and no kinetic parameters were determined, so it bears on the chemistry available at a heme–copper center and not on the rates reported here. Given that the binuclear reaction center is also the site of O₂ reduction to water by Cco, it is possible that the enhancement of Cco/NO activity at low oxygen concentrations is due, in part, to a reduced competition between O₂ and NO₂⁻ for binding to Cco.

A key question left unresolved by prior work is whether Cco/NO is coupled to cellular metabolic state. As O₂ tension falls, mitochondrial oxidative phosphorylation becomes progressively limited and the cellular ADP/ATP ratio rises. This dependence spans the physiological range: in intact cells respiration is nearly independent of oxygen down to about 20 µM, with an apparent Km below 1 µM, yet the cytosolic [ATP]/[ADP] [Pi] ratio varies at all oxygen tensions from air saturation downward, because it is this ratio that adjusts to hold the rate of ATP synthesis constant [27]. The adenine nucleotide signal is therefore available at oxygen concentrations well above those at which the respiratory rate itself is affected. Adenine nucleotides are established allosteric regulators of Cco: ATP inhibits its oxygen-reducing activity (Cco/H₂O), whereas ADP opposes this inhibition [28–30]. The nucleotide and redox state of the cell is a general regulatory input to mitochondrial function, extending from respiratory control to the metabolic reprogramming associated with caloric restriction and aging [31]. If Cco/NO is regulated through an analogous nucleotide-sensitive mechanism, falling O₂ tension and rising ADP/ATP during hypoxia could converge to control local mitochondrial NO formation.

To isolate the Cco-dependent component of nitrite-to-NO conversion and examine its regulation by adenine nucleotides, we used genetically defined yeast strains rather than attempting to reconstitute the complete NO metabolism of a normal eukaryotic cell. Strains YDW29 and YDW27 carry deletions of *YHB1*, which encodes the major yeast NO-consuming flavohemoglobin, and are engineered to express exclusively the Va or Vb isoform of Cco subunit V, respectively. Deletion of *YHB1* removes a dominant NO-consuming activity that would otherwise kinetically mask NO formation at the Cco step. The two genes are switched on or off at very low oxygen concentrations, 0.5–1 µmol l⁻¹ O₂ [32], so a wild-type culture traversing a hypoxic shift changes subunit V composition as it goes. The fixed Va or Vb background cleanly reveals isoform-dependent regulation while diverging from a normal yeast mitochondrion that shifts subunit V composition progressively during the hypoxic transition. Isolated mouse brain mitochondria were used to provide physiological context; brain is one of the two tissues in which steady-state nitrite is highest, owing to its constitutive NOS content [33].

An important experimental constraint is that all Cco/NO assays use exogenous nitrite at 1 mM, a concentration that substantially exceeds steady-state tissue nitrite, which is below 1 µM in most organs and rises to 1.68 ± 0.31 µM in brain and 22.5 ± 9.2 µM in aorta, the two tissues richest in constitutive NOS [13, 33]. These are substrate-driven assays designed to characterize Cco enzymatic capacity under defined O₂, nucleotide, and isoform conditions; they are not measurements at a proven physiological nitrite concentration. Whether Cco/NO operates under local mitochondrial nitrite availability in intact cells depends on the apparent K_m_ of Cco for nitrite, which has not been systematically established.

Here, we show that adenine nucleotides regulate Cco/NO in an oxygen- and isoform-dependent manner. ADP stimulates Cco/NO in yeast and mouse brain mitochondria and shifts its apparent oxygen sensitivity toward lower O₂, whereas the nucleotide-free net rate is not different from zero at any aerobic O₂ tension and increases only at anoxia. Nucleotide regulation is also isoform-dependent: ADP stimulates both Va- and Vb-containing Cco, while ATP slightly inhibits Va-containing Cco but strongly stimulates Vb-containing Cco under anoxic conditions. This reversal of the ATP response suggests that the subunit V isoform context selectively gates the nucleotide response of the nitrite-reductase pathway.

## 2. EXPERIMENTAL PROCEDURES

### 2.1. Yeast Strains, Media, and Growth Conditions

The *Saccharomyces cerevisiae* strains used for this study were: JM43 (*MATα his4-580 trp1-289 leu2-3, 112 ura3-52* [ρ*^+^*]) [34], YDW29 (*MATα his4-580 trp1-289 leu2-3, 112 ura3-52 cox5b::LEU2, yhb1:: Kan^r^* [ρ^+^]) and YDW27 (*MAT*α *his4-580 trp1-289 leu2-3*, *112 ura3-52*, *cox5a*Δ::*URA3*, *COX5b*, *reo1-4*, *yhb1::Kan^r^*, [ρ^+^]) [18]. YDW29 and YDW27 were derived from JM43 and carry deletions of YHB1. In addition, YDW29 carries a deletion in the *COX5b* gene and expresses only the aerobic isoform, Va, of Cco subunit V, while YDW27 carries a deletion in *COX5A* and expresses only the hypoxic isoform, Vb, of Cco subunit V. Yeast cells were grown in SSG-TEA media [35], supplemented with L-histidine, L-tryptophan, L-leucine, uracil, ergosterol (solubilized in Tween 80), and silicone antifoam, as needed [36]. Aerobic cultures were grown on a shaker (200 rpm) at 30 °C and harvested in mid-logarithmic growth phase. For hypoxic shift experiments, a New Brunswick BioFlo 3000 fermentor was used as previously described to bring mid-logarithmic cells from normoxia to anoxia [37]. Cell samples were removed from the fermentor through an ice-cooled coil at the times indicated after anoxia was reached. These samples were washed in ice-cold dH₂O and pelleted at 5,000 rpm for 5 min. Cell pellets were assayed immediately or stored at −80 °C.

### 2.2. Preparation of mitochondria

Yeast mitochondria were prepared from aerobic cultures as described previously [38]. Mouse brain mitochondria were prepared from forebrain tissue of seven-to ten-day-old wild-type mice (reported as C57BL/6 by the animal facility), as previously described [18]. All procedures involving mice were performed in accordance with the guidelines of the Institutional Animal Care and Use Committee of the University of Colorado at Boulder. The protocol number is no longer recoverable, as the laboratory has closed. Animals were euthanized by surgical decapitation, and forebrains from several mice of both sexes were pooled for each preparation. Three preparations were made on separate occasions.

### 2.3. Preparation of Heat Denatured Mitochondria

Purified mitochondria (7 mg of protein/ml) were incubated for 10 min at 60 °C in a medium containing 200 mM potassium phosphate (pH 7.0) and 20 mM dodecyl-β-D-Maltoside. These heat-treated mitochondria exhibited no measurable respiration (data not shown).

### 2.4. Measurement of Cco/NO and oxygen consumption by solubilized mitochondria

Concomitant NO production and O₂ consumption were measured with a 0.7 mm Clark-type NO electrode (Amino-700, Innovative Instruments, Inc, Tampa, FL) and a 2 mm Clark-type O₂ electrode (Oxy-2, Innovative Instruments, Inc, Tampa, FL) respectively, connected to an APOLLO 4000 NO-meter (WPI, Sarasota, Florida) [14]. Except where noted, all solutions were NO₂⁻ free. Measurements were performed at 30 °C using a final reaction volume of 2 mL in a thermostated chamber with a close-fitting lid and fine holes for the electrode and a Hamilton syringe. The Assay Buffer is composed of 0.5 mM dodecyl-β-D-Maltoside in 200 mM Tris-Cl (pH 7.0). Assays generally used 50 µg of mitochondrial protein, which was solubilized in Assay Buffer for 10 min prior to the addition of: cytochrome c to a final concentration of 10 µM; ascorbate (pH 7.0) to a final concentration of 0.5 mM; and N,N,N′,N′-tetramethyl-p-phenylenediamine dihydrochloride (TMPD) to a final concentration of 0.5 mM. Then, the lid on the chamber was closed and additional components (e.g., NO₂⁻, ATP, ADP) were added, as needed, through the lid port using a Hamilton syringe. Detergent solubilization is required by the design of this assay rather than adopted for convenience. Electrons are delivered to Cco from externally added cytochrome c reduced by ascorbate/TMPD, an interaction whose kinetics have been characterized for the isoforms of yeast cytochrome c and of Cco subunit V [39]; in mitochondria with an intact outer membrane exogenous cytochrome c has no access to its binding site on Cco, and solubilization is what makes the electron-donor system operative. Disrupting the inner membrane also abolishes the proton gradient and the membrane potential, so the preparation is uncoupled by construction: a respiratory control ratio is not defined for it, and any effect of adenine nucleotides on either activity must act on the enzyme itself rather than through the proton-motive force. Solubilization also gives externally added adenine nucleotides direct access to the enzyme, so that the nucleotide dependence measured here is a property of Cco rather than of nucleotide transport across two membranes. The consequence is that all activities reported below are catalytic capacities of the enzyme in a defined medium, not fluxes in an energetically coupled organelle. Solubilization was complete under these conditions: addition of the mitochondrial suspension to Assay Buffer abolished its turbidity, giving an optically clear solution. Functional confirmation that exogenous cytochrome c reaches its site on Cco is given in Figure 1C, where added cytochrome c supports NO synthesis with an apparent Km of 0.5–1.0 µM and saturation at 10 µM, and where its omission reduces the rate to less than one third.

**Figure 1.**
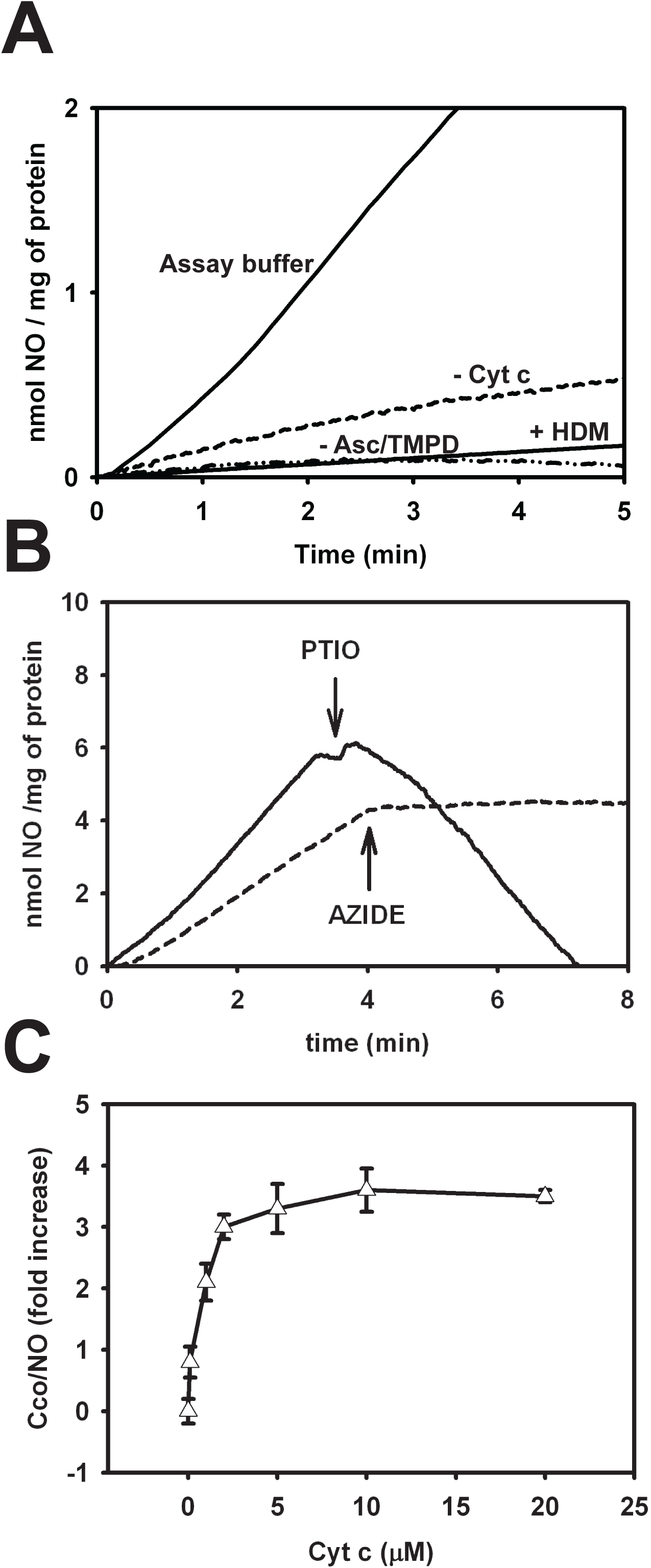
Cco/NO activity in yeast detergent-solubilized mitochondria. Mitochondria were from strain YDW29, which expresses only the Va isoform of Cco subunit V and lacks YHb; assays were under anoxia (buffer pre-bubbled with N₂) except where oxygen is varied. (A) Polarographic measurement of NO production in complete assay buffer, in assay buffer lacking cytochrome c, in assay buffer lacking ascorbate/TMPD, and with heat denatured mitochondria (HDM). Solubilized yeast mitochondria were assayed for NO production in assay buffer pre-bubbled with nitrogen for 10 min in the presence of 1 mM NO₂⁻. The chamber was closed and NO synthesis measured with an NO electrode. (B) Effects of PTIO (2-phenyl-4,4,5,5-tetramethylimidazoline-3-oxide-1-oxyl) and azide on NO synthesis in complete assay buffer. The ordinate is the free NO concentration measured by the electrode, not the rate of NO synthesis. At the times indicated by arrows 1 mM PTIO (solid line) or 5 mM sodium azide (dashed line) was added: PTIO lowers the signal by scavenging the NO already present, whereas azide, a Cco inhibitor, halts further NO synthesis so that the signal plateaus. (C) Effect of added cytochrome c on the initial rate of NO synthesis. Mean and standard deviation values in C are for three independent measurements.

### 2.5. Measurement of Cco/NO activity in the presence of adenine nucleotides

The ADP and ATP effects on Cco/NO and oxygen consumption were measured as indicated above with the addition of different concentrations of the adenine nucleotides. ATP concentration was kept constant by the use of an ATP-regenerating system: 10 mM phosphoenolpyruvate (PEP), 10 units/ml of pyruvate kinase (PK), 5 mM MgSO₄ and different concentrations of MgATP, as indicated.

### 2.6. ATP/ADP determination

Adenine nucleotides were extracted from whole cell yeast with 50 mAU/mL proteinase K (GibcoBRL #25530-015) as previously described [40]. Cellular ATP and ADP levels were quantified from these extracts using ATP bioluminescent Assay Kit (Sigma FLAA). ATP was quantified directly and immediately as described in the kit. Briefly, an aliquot of the extracted sample was mixed with the kit’s Sigma Assay Mix (FL-AAM) and the emission at 500 nm was measured (Molecular Devices SpectrMax M5). ADP was quantified indirectly by the following method: First, the extracted ATP was removed from the assay by conversion to AMP using ATP sulfurylase (Sigma A8957) according to the manufacturer’s instructions (50 mM Tris-HCl (pH 8.0), 5 mM MgCl₂, 10 mM Na₂MoO₄, 2.5 mM GMP, and 30 µg/mL ATP sulfurylase). Sulfurylase was then heat-inactivated at 95 °C and the baseline ATP levels were measured. Next, the extracted ADP was converted to ATP using the ATP-regenerating system described above. The resulting ATP levels were then measured and the baseline ATP was subtracted to achieve the cellular ADP level measurements. Because ATP is enzymatically removed before this step, the increment measured reports nucleotide converted by pyruvate kinase, which acts on ADP and, less efficiently, on other nucleoside diphosphates. A broader quantity is measured by total nucleoside diphosphate determinations using 3¹P-NMR [41].

### 2.7. Miscellaneous methods

Protein concentration was determined by the Bradford assay (Pierce Biotechnology #23238) with BSA as a standard. Cytochrome c oxidase from *S. cerevisiae* was prepared by Method 1 described previously [42].

### 2.8. Data analysis

Values are means ± standard deviation of three independent measurements, each made on a separate mitochondrial preparation (Tables I and III), or means ± standard error of the individual recordings, with n given in the legend (Table II). Instantaneous rates of NO production and of oxygen consumption were obtained from the slope of the corresponding electrode trace at each indicated oxygen concentration. Group comparisons were made by two-sided Welch’s t-test, computed from the reported means, standard deviations and n for Tables I and III and from the individual recordings for Table II, with Holm correction within each family of related comparisons; the resulting t, degrees of freedom and adjusted p values are reported in the text and in the point-by-point response to the reviewers. Net rates whose mean did not differ from zero (one-sample t-test, P > 0.05) are shown as n.d. in Table II, and the individual recordings are given in Supplementary Table S1. Because the O₂ points of a recording are successive readings from the same trace, the oxygen dependence was tested with one value per recording, the slope of the rate against ln[O₂] over the aerobic range, compared between conditions by Welch’s t-test. The trends shown in Figures 2–6 are described qualitatively, and differences smaller than the spread of the nucleotide-free control are not interpreted.

**Table I.**
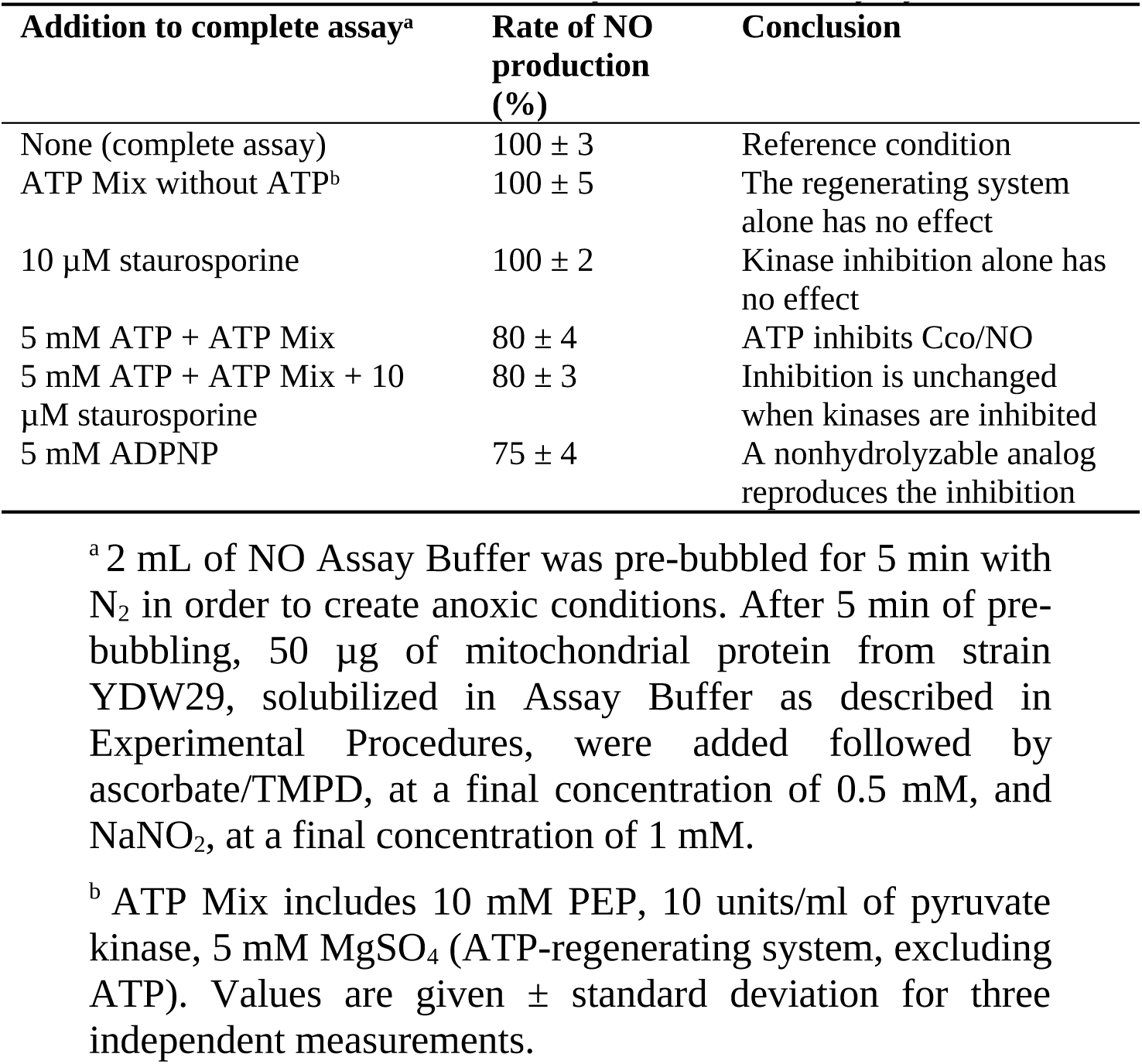
Immediate inhibition of Cco/NO activity by ATP.

**Table II.** Oxygen sensitivity of brain Cco/NO in the presence of ADP and ATP.

|  | NO production (nmol NO·mg protein<sup>−1</sup>·min<sup>−1</sup>)<sup>a</sup> |  |  |
|---|---|---|---|
| **O<sub>2</sub> (µM)** | **Control** | **+ ADP** | **+ ATP** |
| 175 | — | 0.040 ± 0.005 | 0.034 ± 0.001 |
| 150 | — | 0.050 ± 0.003 | n.d. |
| 100 | n.d. | 0.066 ± 0.006 | n.d. |
| 75 | n.d. | 0.082 ± 0.007 | n.d. |
| 50 | n.d. | 0.120 ± 0.009 | n.d. |
| 40 | n.d. | 0.147 ± 0.010 | n.d. |
| 30 | n.d. | 0.191 ± 0.014 | n.d. |
| 20 | n.d. | 0.268 ± 0.023 | n.d. |
| 15 | n.d. | 0.333 ± 0.031 | n.d. |
| 10 | n.d. | 0.443 ± 0.050 | n.d. |
| 5 | n.d. | 0.669 ± 0.104 | n.d. |
| 0 | 0.304 ± 0.018 | 1.209 ± 0.124 | 0.237 ± 0.014 |

<sup>a</sup> Mouse brain mitochondria, solubilized in Assay Buffer as described in Experimental Procedures, prepared from pooled forebrains of several mice of both sexes, were assayed for NO production pre-bubbled with air in the presence of 1 mM NO<sub>2</sub><sup>−</sup> and 0.5 mM ascorbate/TMPD. NO production was measured with an NO electrode, and dissolved oxygen concentration was measured concomitantly with an O<sub>2</sub> electrode in a closed chamber. The activities were measured in the absence of any externally added adenine nucleotide or in the presence of 5 mM ADP or 5 mM ATP + ATP Mix. NO production was normalized to total mitochondrial protein. Values are means ± standard error of the individual recordings: n = 4 with ADP and with ATP (n = 2 with ATP at 150 and 175 µM O<sub>2</sub>) and n = 2 without nucleotide (n = 1 at 150 and 175 µM O<sub>2</sub>); with ADP and with ATP, the four recordings were made two on each of two days. Rates are slopes of the NO-electrode trace and are therefore net rates (formation minus consumption). n.d., net rate not different from zero (one-sample t-test, P > 0.05); —, single recording, no dispersion (value in Supplementary Table S1). The individual recordings are given in Supplementary Table S1.

**Table III.**
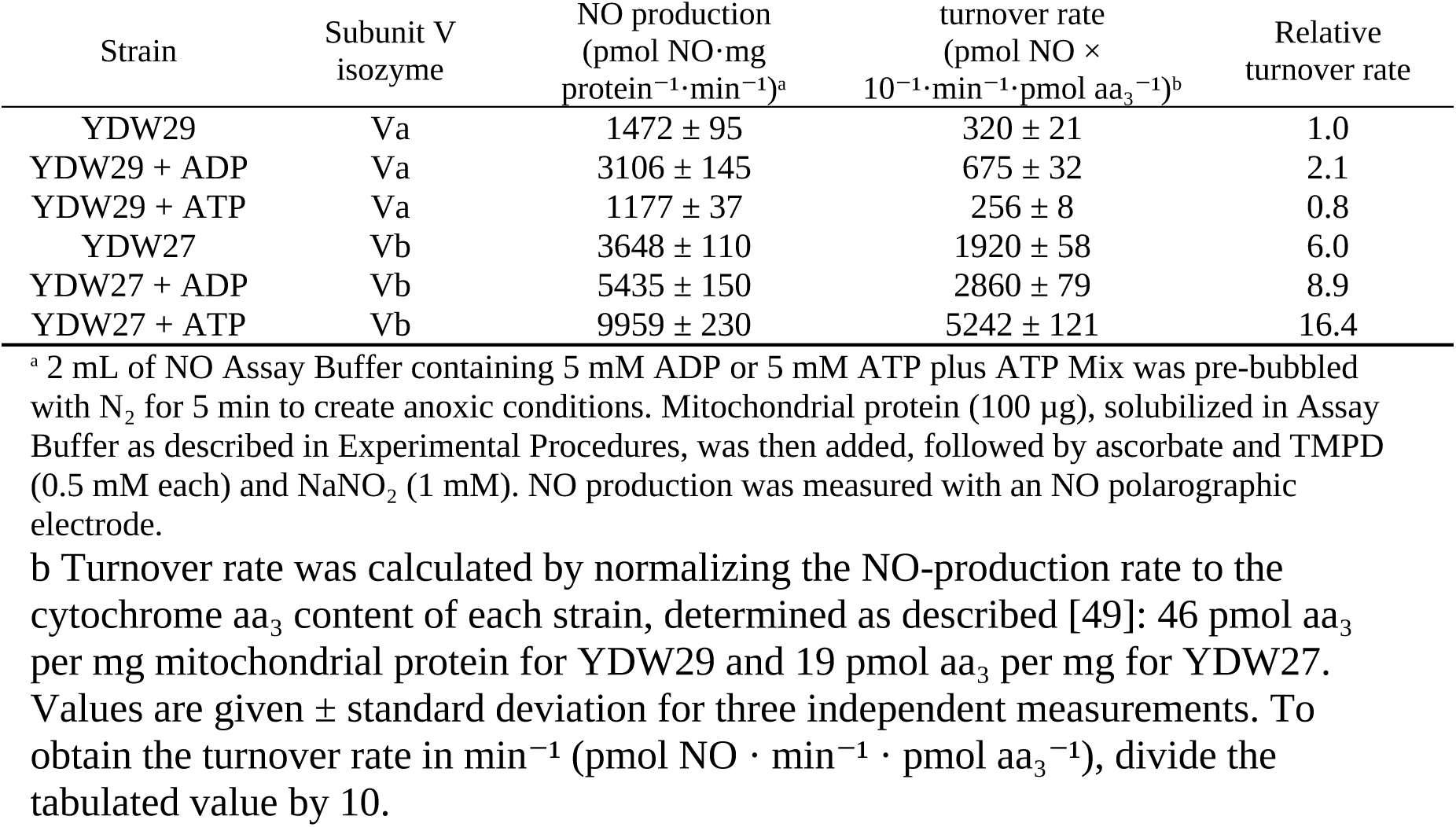
ADP and ATP modulation of nitrite-dependent NO production by Cco containing different subunit V isoforms.

| Strain | Subunit V<br>isozyme | NO production<br>(pmol NO·mg<br>protein<sup>−1</sup>·min<sup>−1</sup>)<sup>a</sup> | turnover rate<br>(pmol NO ×<br>10<sup>−1</sup>·min<sup>−1</sup>·pmol aa<sub>3</sub><sup>−1</sup>)<sup>b</sup> | Relative<br>turnover rate |
|---|---|---|---|---|
| YDW29 | Va | 1472 ± 95 | 320 ± 21 | 1.0 |
| YDW29 + ADP | Va | 3106 ± 145 | 675 ± 32 | 2.1 |
| YDW29 + ATP | Va | 1177 ± 37 | 256 ± 8 | 0.8 |
| YDW27 | Vb | 3648 ± 110 | 1920 ± 58 | 6.0 |
| YDW27 + ADP | Vb | 5435 ± 150 | 2860 ± 79 | 8.9 |
| YDW27 + ATP | Vb | 9959 ± 230 | 5242 ± 121 | 16.4 |

<sup>a</sup> 2 mL of NO Assay Buffer containing 5 mM ADP or 5 mM ATP plus ATP Mix was pre-bubbled with N<sub>2</sub> for 5 min to create anoxic conditions. Mitochondrial protein (100 µg), solubilized in Assay Buffer as described in Experimental Procedures, was then added, followed by ascorbate and TMPD (0.5 mM each) and NaNO<sub>2</sub> (1 mM). NO production was measured with an NO polarographic electrode.
b Turnover rate was calculated by normalizing the NO-production rate to the cytochrome aa<sub>3</sub> content of each strain, determined as described [49]: 46 pmol aa<sub>3</sub> per mg mitochondrial protein for YDW29 and 19 pmol aa<sub>3</sub> per mg for YDW27. Values are given ± standard deviation for three independent measurements. To obtain the turnover rate in min<sup>−1</sup> (pmol NO · min<sup>−1</sup> · pmol aa<sub>3</sub><sup>−1</sup>), divide the tabulated value by 10.

**Figure 2.**
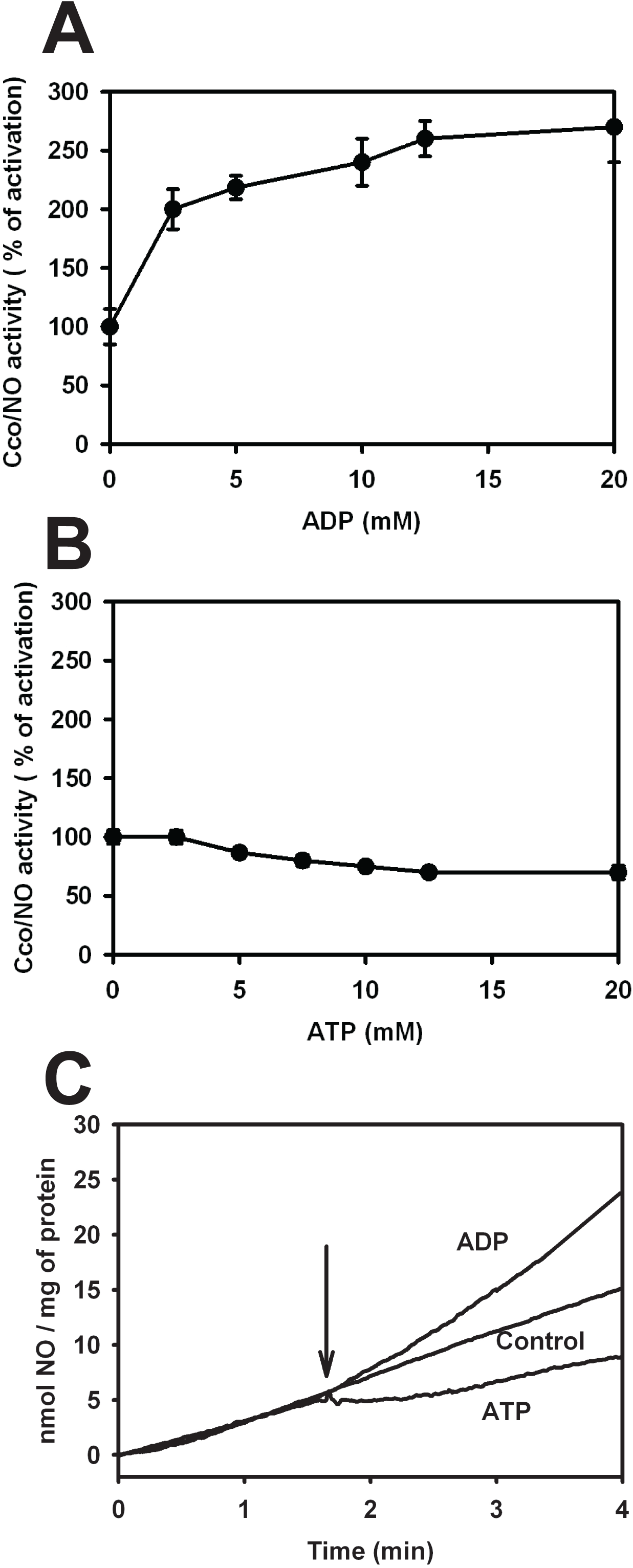
Modulation of Cco/NO activity in yeast detergent-solubilized mitochondria by adenine nucleotides. Mitochondria were from strain YDW29 (Va isoform), assayed under anoxia. NO synthesis by holo-cytochrome c oxidase carrying the subunit Va isoform was determined polarographically with an NO electrode as indicated in the Legend to Figure 1. The plots show the percent change of Cco/NO activity (rate of Cco/NO-catalyzed NO synthesis) in the presence of different concentrations of ADP (A) or ATP in an ATP-regenerating system (B). Cco/NO activities (100%) were normalized to the rate of Cco/NO-catalyzed NO synthesis (0.5 nmol NO·mg protein⁻¹·min⁻¹) in the absence of added adenine nucleotides, so that values below 100% indicate inhibition. (C) Addition of ATP or ADP has an immediate effect on Cco/NO activity. NO synthesis was initiated in the absence of added adenine nucleotide and, at the time indicated by the arrow, 5 mM ADP or 5 mM ATP (with the ATP-regenerating system) was added to the chamber and NO levels were measured. Mean and standard deviation values in A and B are for three independent measurements.

**Figure 3.**
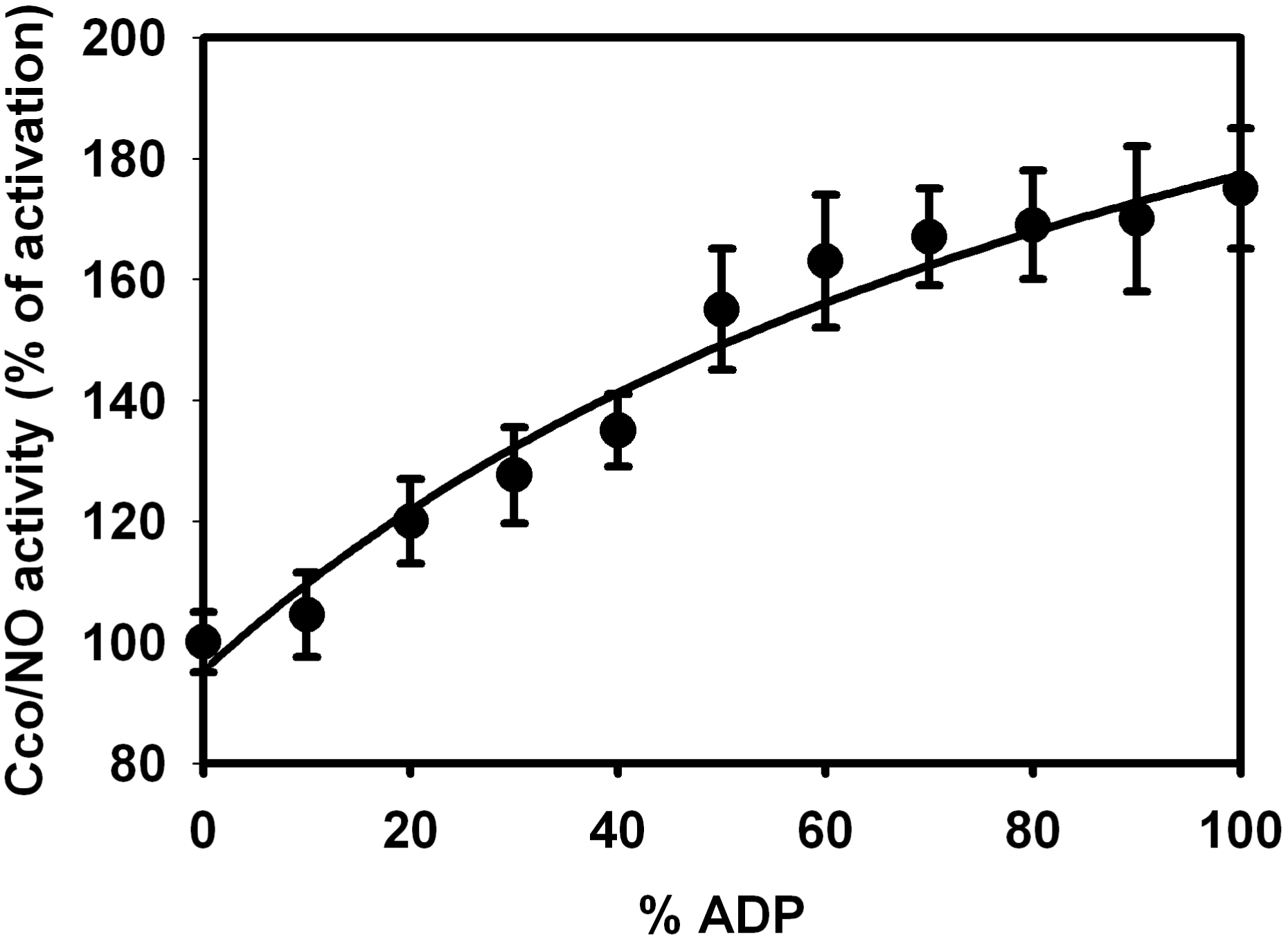
The Cco/NO activity of purified yeast Cco is affected by ADP/ATP ratio. The enzyme was purified from aerobically grown JM43 cells, in which Va is the only subunit V isoform detectable by immunoblot [59]; assays were under anoxia. Purified yeast Cco was suspended in Assay Buffer and NO activity was measured over the range from 0 to 100% ADP (where total added adenine nucleotides [ATP + ADP] = 10 mM), as described in the Legend to Figure 1. Cco/NO activities (100%) were normalized to the rate at 0% ADP, that is with 10 mM ATP; in Figures 2 and 4 the 100% value is instead the rate in the absence of added adenine nucleotides. Mean and standard deviation values are for three independent measurements.

**Figure 4.**
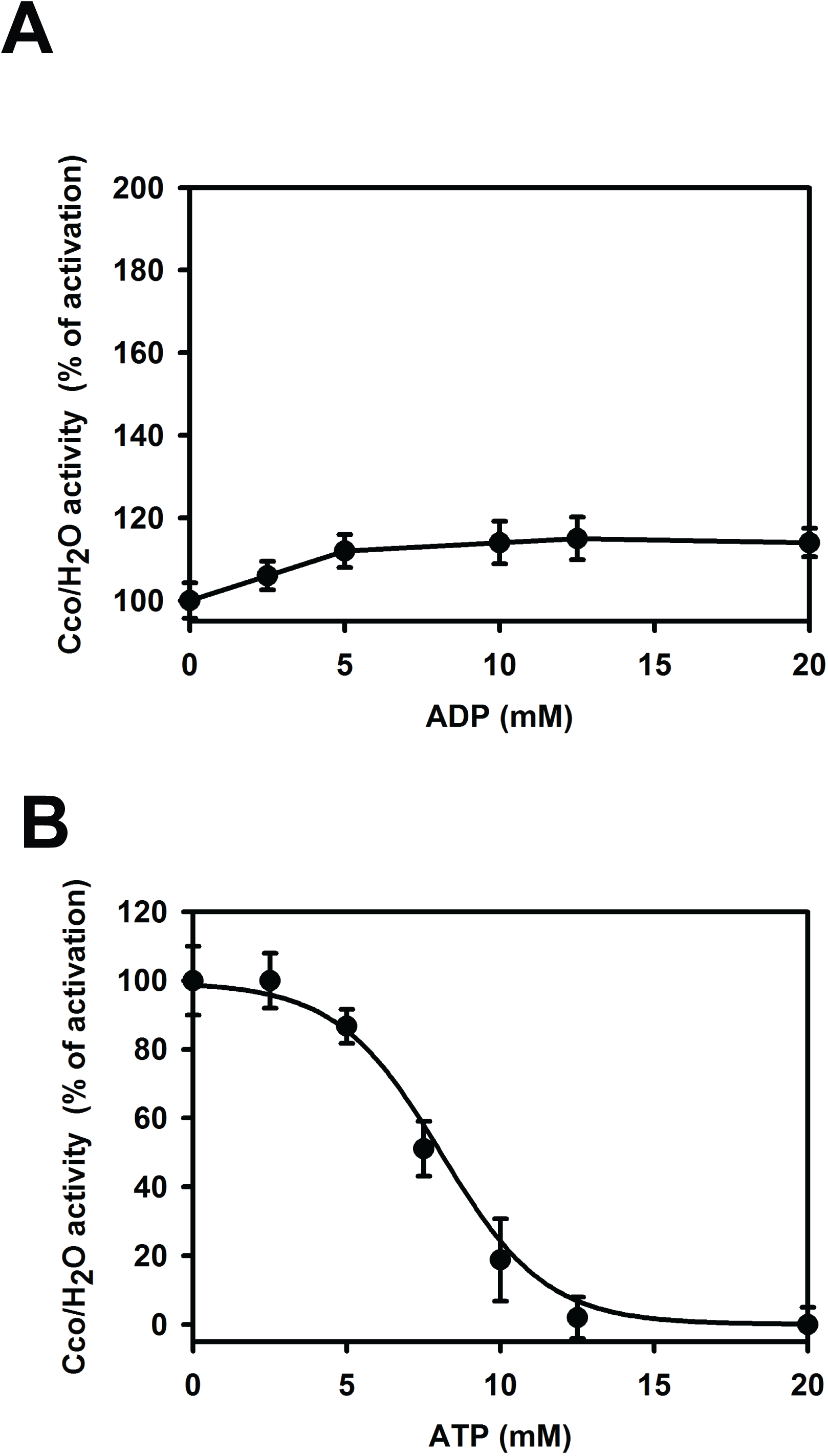
Effects of ADP and ATP on the rate of oxygen consumption by yeast holo-cytochrome c oxidase containing the subunit Va isoform. Oxygen consumption (Cco/H₂O activity) in detergent-solubilized YDW29 mitochondria was measured without added nitrite using an oxygen electrode after 10 min equilibration with air. Plots show percent change in Cco/H₂O activity with ADP (A) or ATP plus the ATP-regenerating system (B). Activities were normalized to the nucleotide-free rate of 0.95 µmol O₂·mg protein⁻¹·min⁻¹. Mean and standard deviation values are for three independent measurements.

**Figure 5.**
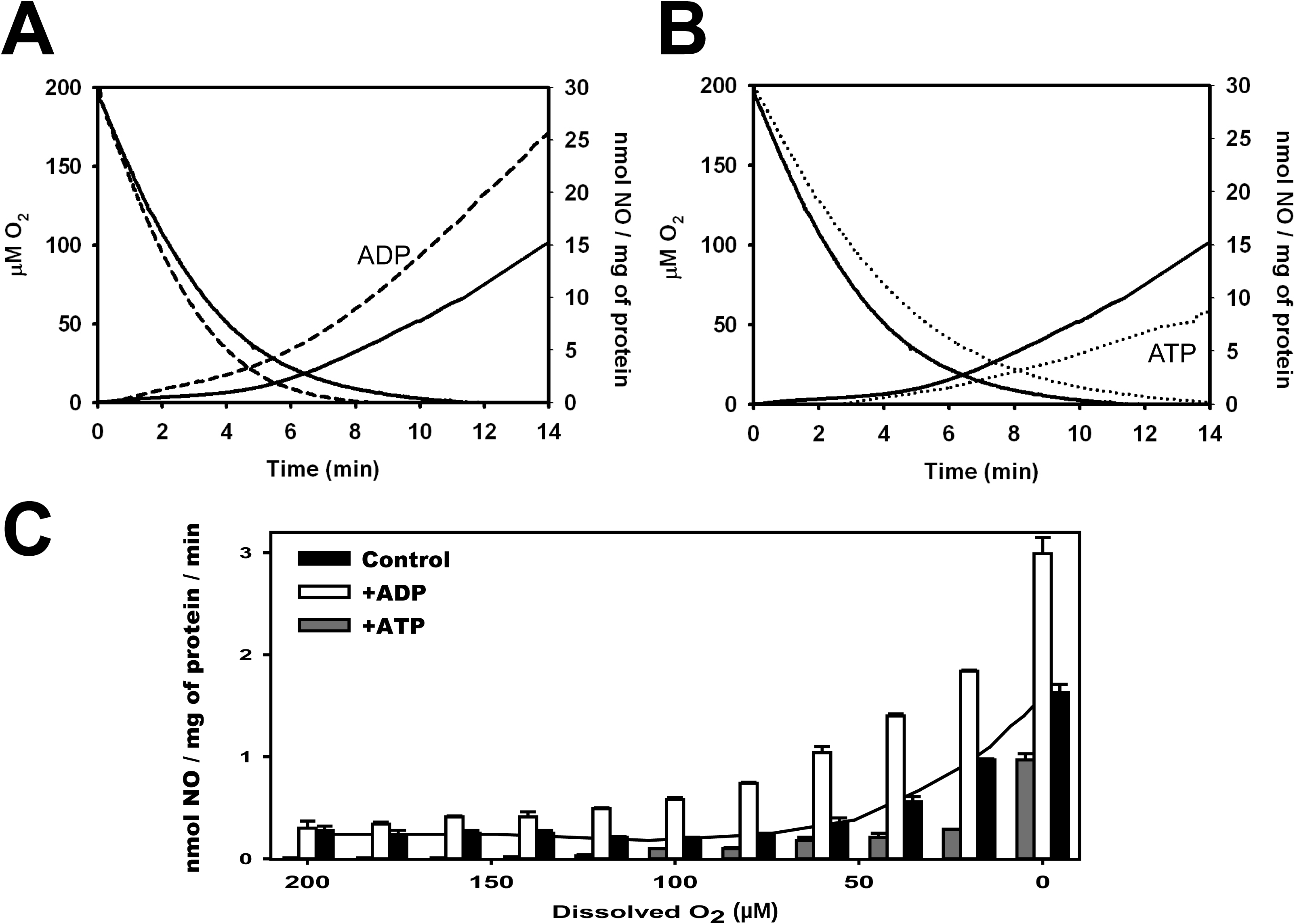
Adenine nucleotides affect the oxygen sensitivity of yeast Cco/NO. Mitochondria were from strain YDW29 (Va isoform). Detergent-solubilized yeast mitochondria were assayed simultaneously for oxygen consumption and Cco/NO activity in air-equilibrated assay buffer containing 1 mM nitrite. Activities were measured without nucleotide (solid lines) and with 5 mM ADP (A, long-dashed lines) or 5 mM ATP plus the ATP-regenerating system (B, short-dotted lines). In A and B oxygen falls with time and NO accumulates. Instantaneous Cco/NO rates at the indicated oxygen concentrations are shown in C; no added nucleotide, black bars; 5 mM ADP, open bars; 5 mM ATP plus the ATP-regenerating system, grey bars. Oxygen decreases from left to right, following the time course in A and B. In C the line represents the fit for the Control data. Mean and standard deviation values in C are for three independent measurements.

**Figure 6.**
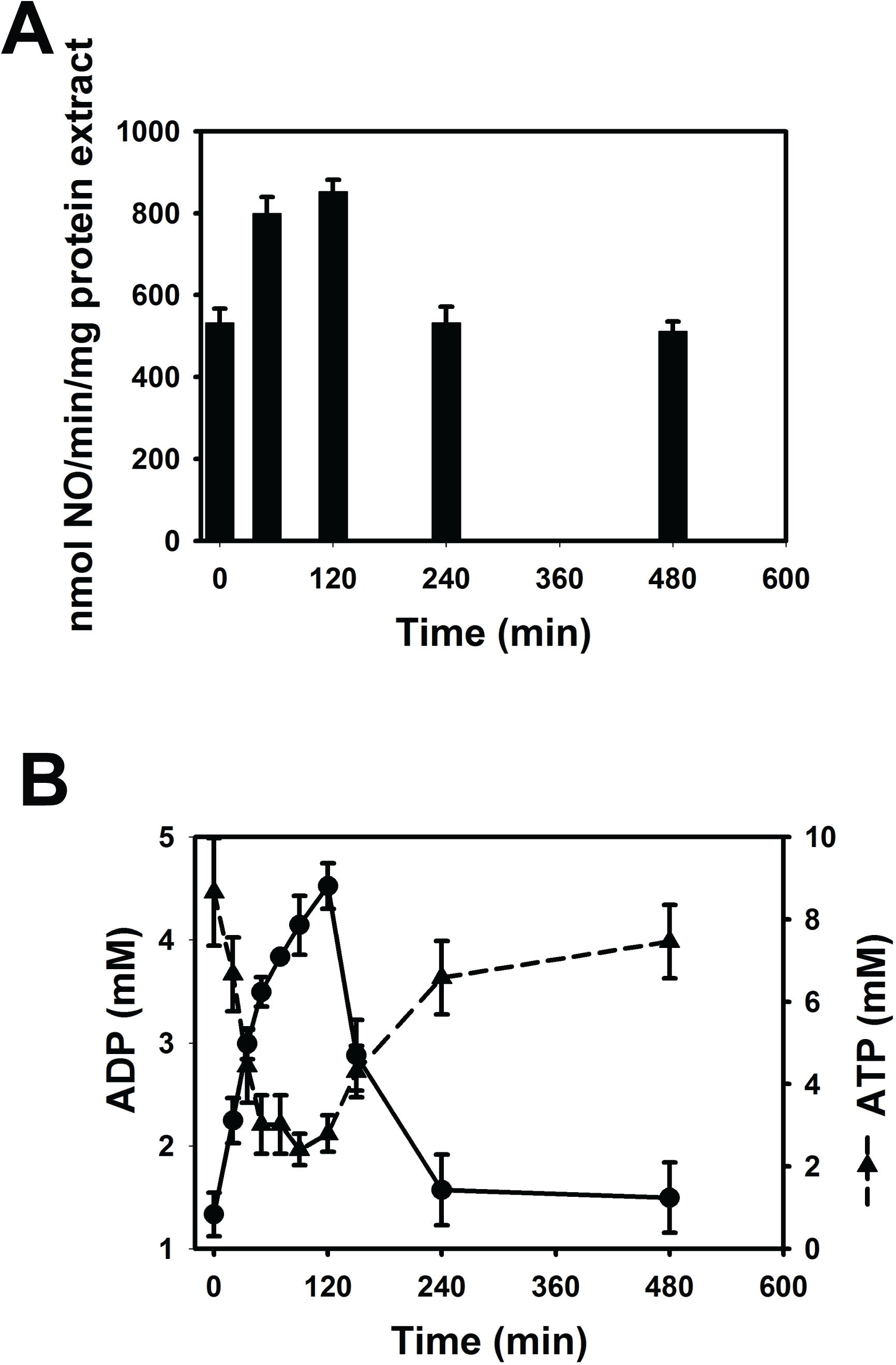
Cco/NO capacity and adenine nucleotide levels increase transiently during a hypoxic shift. JM43 yeast cells were maintained in steady-state normoxic growth in a fermentor sparged with air for six generations. The process gas was shifted to 97.5% N₂ and 2.5% CO₂. Cells were sampled at the indicated times for nucleotide determination (B) and subsequent assay of Cco/NO activity in mitochondrial preparations (A). In B, ADP is shown on the left axis and ATP on the right axis, so that both components of the ADP/ATP ratio can be read directly. The Cco/NO measurements are not direct intact-cell NO measurements. ADP was determined after enzymatic removal of ATP, by conversion with pyruvate kinase (see Experimental Procedures). Mean and standard deviation values are for three independent measurements.

## 3. RESULTS AND DISCUSSION

### 3.1. ATP and ADP Have Differential Effects on Cco/NO activity

To evaluate the effects of adenine nucleotides on Cco/NO we used an assay with detergent-solubilized yeast mitochondria, externally added cytochrome c, ascorbate, TMPD, and NO₂⁻ [18]. Because this assay system uses detergent-solubilized mitochondria, it allows Cco to be in contact with externally added adenine nucleotides and cytochrome c. Moreover, because the electron donor pair TMPD/ascorbate feeds electrons directly to cytochrome c (both free and Cco-bound), the assay isolates the production of NO to the terminal portion of the respiratory chain. In order to control for any subunit V isoform-specific effects, our initial experiments were done with mitochondria from yeast strain YDW29, which expresses only the Va isoform of Cco subunit V and which is deleted for the YHb hemoglobin, an NO dioxygenase which consumes NO in yeast [43]. From Figure 1A, it is clear that optimal NO synthesis requires functional mitochondria, ascorbate, TMPD, and cytochrome c. NO synthesis is not observed with heat denatured mitochondria or in the absence of ascorbate or TMPD. Omission of added cytochrome c reduces the rate of NO synthesis to less than a third of that supported by the complete assay system. As reported earlier [18], the reduced level of NO synthesis observed in the absence of added cytochrome c is supported by endogenous cytochrome c. Addition of the NO scavenger, PTIO (2-Phenyl-4,4,5,5-tetramethylimidazoline-3-oxide-1-oxyl), immediately reduced the observed NO signal, and NO synthesis is immediately inhibited by 5 mM sodium azide, a Cco inhibitor (Figure 1B).

Added cytochrome c increases the rate of NO synthesis in a hyperbolic fashion (Figure 1C), with an apparent Km of 0.5–1.0 µM and maximal activity at 10 µM, the concentration used in subsequent assays to maintain cytochrome c in excess. We next tested a range of added ADP and ATP concentrations as biochemical perturbations of Cco/NO activity. Increasing ADP up to 20 mM stimulated Cco/NO, whereas increasing ATP inhibited the Va-containing enzyme (Figures 2A and 2B). These externally added concentrations in detergent-solubilized mitochondria define an assay response and should not be equated directly with free matrix nucleotide concentrations in intact cells. The response to either nucleotide was immediate (Figure 2C), supporting a direct allosteric effect rather than a delayed post-translational modification. Staurosporine, a nonspecific protein kinase inhibitor [44], did not markedly alter ATP inhibition, and the nonhydrolyzable analog ADPNP produced an inhibitory effect similar to ATP (Table I). Both the ATP and the ADPNP inhibition were significant relative to the complete assay (Welch’s t-test, Holm-adjusted p = 0.012 and 0.007, respectively), whereas the ATP-regenerating system without ATP and staurosporine alone had no effect (adjusted p = 1.0). These findings argue that ATP hydrolysis and protein phosphorylation are not required for the observed inhibition.

Because ATP and ADP coexist at concentrations that vary with cellular and mitochondrial energetic state, we next examined Cco/NO activity in the presence of both nucleotides. For this study, we used highly purified yeast cytochrome c oxidase. As expected, NO synthesis by purified Cco responds to adenine nucleotides, just as it does in the solubilized mitochondria used above, and increases with increasing ADP:ATP ratios (Figure 3).

This is the first report that ADP and ATP have differential effects on Cco/NO activity. Their effects on the Va-containing nitrite-reductase pathway differ in magnitude from their effects on oxygen reduction to water (Cco/H₂O) [28, 29, 45, 46]. ADP markedly stimulates Cco/NO while only modestly increasing Cco/H₂O, whereas ATP inhibits both Va activities and has a much larger effect on oxygen consumption (Figure 4A and 4B). The difference is quantitative. At 12.5 mM ADP, Cco/NO rises to approximately 260% of the nucleotide-free rate (Figure 2A) whereas Cco/H₂O rises only to approximately 115% (Figure 4A). With ATP the two activities track together up to 5 mM, where both retain approximately 88% of the nucleotide-free rate, and diverge sharply above it: at 12.5 mM ATP, Cco/H₂O is essentially abolished (Figure 4B) while Cco/NO retains approximately 70% of its activity (Figure 2B). Thus, adenine nucleotides do not simply scale both catalytic outputs in parallel, and at the ATP concentrations that suppress oxygen reduction most of the nitrite-reductase activity is preserved.

In intact yeast mitochondria the same nucleotides act differently, and the contrast is informative. Removal of ADP lowered nitrite-dependent NO production to 12% of control, and either ATP or the uncoupler dinitrophenol restored it, which led to the conclusion that ADP acts there by increasing the rate of electron transport through the respiratory chain [14]. That route requires a proton gradient across an intact inner membrane and is therefore unavailable in the solubilized preparation used here, which has neither a proton gradient nor a functional F₁F₀ ATP synthase. The opposite sign of the ATP effect in the two preparations is thus expected rather than contradictory: in a coupled organelle the dominant effect of adenine nucleotides is on electron delivery, whereas the effects reported here are properties of the enzyme.

### 3.2. Adenine nucleotides alter the oxygen sensitivity of Cco/NO

The finding that ADP stimulates Cco/NO activity led us to wonder if ADP also alters the oxygen sensitivity of the Cco/NO reaction. To clarify the effect of oxygen concentration *per se* on the rate of Cco/NO we calculated the instantaneous rate of NO synthesis at different oxygen concentrations in the presence and absence of ADP or ATP. Figures 5A and 5B show the simultaneous consumption of oxygen by Cco (Cco/H₂O) and production of NO by Cco/NO, in the absence and presence of ADP or ATP. Added ADP increases the rate of NO production (Cco/NO activity) and slightly increases the rate of oxygen consumption (Cco/H₂O activity). In contrast, ATP decreases the rate of NO production and oxygen consumption. As shown in Figure 5C, ADP decreases the apparent inhibition of Cco/NO by oxygen, allowing NO synthesis at much higher oxygen concentrations than previously reported [14]. Indeed, in the presence of added ADP, NO is produced at dissolved O₂ concentrations up to 120–140 µM O₂. Conversely, ATP enhances the oxygen inhibition of Cco/NO activity even at very low O₂ levels.

The yeast experiments above resolve the oxygen dependence up to the point at which the chamber is depleted. To extend the comparison to a mammalian preparation, and to cover the oxygen range of vertebrate tissue in a single series, we repeated the measurement in isolated mouse brain mitochondria. We measured instantaneous NO-production rates at twelve dissolved O₂ concentrations from 175 µM to anoxia, without added nucleotide, with 5 mM ADP, or with 5 mM ATP maintained by an ATP-regenerating system (Table II). Without added nucleotide, the net rate was not different from zero at any aerobic O₂ tension and increased to 0.304 ± 0.018 nmol NO·mg protein⁻¹·min⁻¹ (mean ± SE, n = 2) at anoxia.

With 5 mM ADP, the mean rate increased progressively as O₂ fell, reaching 1.209 ± 0.124 nmol NO·mg protein⁻¹·min⁻¹ at anoxia (n = 4), approximately 4.0-fold above the nucleotide-free mean at anoxia. With one slope per recording (rate against ln[O₂] over the aerobic range), the oxygen dependence with ADP (−0.164 ± 0.025 nmol NO·mg protein⁻¹·min⁻¹ per ln unit, n = 4) differed from that of the nucleotide-free control (+0.006 ± 0.010, n = 2; Welch’s t-test, p = 0.004, or p = 0.027 with the two ADP recordings of each day averaged), consistent with a change in the oxygen dependence rather than a uniform scaling of the baseline rate. The O₂ concentrations at which the ADP effect is significant (10 µM and below; see below) lie below the approximately 25 µM of muscularized capillaries and below the P50 of human hemoglobin, 27 Torr (approximately 36 µM) [10, 47].

With 5 mM ATP, the net rate was not different from the nucleotide-free control at any O₂ tension, including anoxia (0.237 ± 0.014 nmol NO·mg protein⁻¹·min⁻¹, n = 4). Welch’s t-tests on the individual recordings (Supplementary Table S1) show that ADP increased the Cco/NO rate relative to the nucleotide-free control at low O₂, with the difference significant at 10 and 5 µM O₂ and at anoxia after Holm correction and not resolved at 15 µM O₂ and above, whereas the ATP condition did not differ from the control at any O₂ tension; the ADP and ATP rates differed at every O₂ tension up to 100 µM; the test statistics follow from Supplementary Table S1. These results show that the robust effect of ADP in brain mitochondria is a shift of the apparent oxygen sensitivity of Cco/NO toward low O₂.

Together, these results show that the apparent oxygen sensitivity of Cco/NO depends on the nucleotide environment. ADP therefore reduces the apparent oxygen inhibition of Cco/NO under these assay conditions. The mechanism remains unclear, although the absence of ADP-mediated inhibition of oxygen consumption argues against simple obstruction of O₂ access or reduction at the binuclear center. One point deserves to be stated explicitly, because it is easily mistaken: nitrite at 1 mM does not inhibit the oxidase at the oxygen tensions over which the oxidase is turning over. In Figure 5 the oxygen present at air equilibration is consumed to exhaustion with 1 mM nitrite in the chamber throughout, and NO formation begins only once the dissolved oxygen concentration has fallen below about 20 µM, as reported previously [14]. This is consistent with, rather than contrary to, the established inhibition of Cco by NO, which is competitive with oxygen and therefore weak at the tensions over which the oxygen trace is recorded [48]. The two processes are sequential along the same trace: Cco reduces nitrite as oxygen becomes limiting, and the NO so formed can then act on the enzyme that produced it. The two activities can also be dissociated experimentally, since low-intensity light stimulates Cco/NO without affecting Cco/H₂O [18]. Under these conditions nitrite behaves as a substrate of Cco rather than as a respiratory poison.

### 3.3. ATP and ADP differentially regulate Cco/NO catalyzed by yeast Cco isozymes containing different subunit V isoforms

To test isoform dependence, we used mitochondria from YDW29 and YDW27, which lack YHb and express exclusively Va or Vb, respectively. NO-production rates were normalized to cytochrome aa₃ content as described by Waterland et al. [49] and measured in anoxia, where Cco/NO activity was maximal. Table III shows that the Vb enzyme has a higher turnover rate than the Va enzyme in this assay. A comparable isoform difference is seen in the oxidase reaction, where Vb raises the catalytic constant three-to fourfold relative to Va by altering an internal electron-transfer step between heme a and the heme a₃–Cu_B_ binuclear center [32]. Because nitrite is reduced at that same binuclear center, a change in the rate at which electrons are delivered to it offers a parsimonious account of why both activities are higher in the Vb enzyme. The same ranking holds in intact cells: in vivo turnover rates for nitrite-dependent NO production, obtained in anoxic whole yeast cells and normalized to intracellular cytochrome aa₃, are 4.5-to 7-fold higher for Vb-containing than for Va-containing Cco [16]. The isoform effect is therefore not an artefact of the solubilized preparation. ADP stimulates both forms. ATP slightly inhibits Va but increases Vb activity 2.7-fold relative to Vb baseline; the value of 16.4-fold compares Vb plus ATP with Va baseline and is therefore a cross-isoform, not a within-Vb, comparison. Each of these nucleotide effects, and the Vb-versus-Va difference, is significant (Welch’s t-test, Holm-adjusted p ≤ 0.023; Table III), and the opposite sign of the ATP effect in the two isoforms is itself highly significant (interaction test, p < 0.001). These findings identify an isoform-dependent reversal of the ATP response, most simply explained by distinct nucleotide-sensitive environments in Va- and Vb-containing Cco. Site-directed mutagenesis and binding studies will be required to locate the responsible site.

### 3.4. Catalytic capacity of Cco/NO relative to other NO-producing systems

Table III permits an enzyme-normalized test of whether Cco/NO represents a single stoichiometric reduction or sustained catalysis. Applying the 10⁻¹ factor specified in the reported turnover unit gives calculated rates of 0.53, 1.13, and 0.43 NO molecules·Cco⁻¹·s⁻¹ for Va baseline, Va plus ADP, and Va plus ATP, respectively, and 3.20, 4.77, and 8.74 NO molecules·Cco⁻¹·s⁻¹ for the corresponding Vb conditions. Thus, in the anoxic assay with 1 mM nitrite, added cytochrome c, ascorbate, and TMPD, Cco is repeatedly reduced and supports multi-turnover nitrite reduction rather than a single terminal heme reaction. These values are turnover rates observed at a single nitrite concentration, 1 mM, and are not catalytic constants. Nitrite was not titrated in this system and saturation was not demonstrated, so they are lower bounds on the turnover the enzyme can sustain; no kcat, apparent Km or second-order rate constant can be derived from the present data. Obtaining those parameters would require a nitrite titration in the solubilized system, which was not performed.

For scale, Rodríguez-Crespo et al. reported a specific activity of 95–120 nmol·min⁻¹·mg⁻¹ for purified eNOS, measured by both NO and citrulline production [50], Harteneck et al. reported 1.2 ± 0.04 µmol·min⁻¹·mg⁻¹ for purified nNOS [51], and Feng et al. reported 1100–1300 nmol·min⁻¹·mg⁻¹ for purified iNOS [52]. Using approximate monomer masses of 135, 160, and 130 kDa, respectively, these activities correspond to about 0.21–0.27 s⁻¹ for eNOS, 3.2 s⁻¹ for nNOS, and 2.4–2.8 s⁻¹ for iNOS. The turnover rates measured here therefore fall in the same broad order as those of the purified NOS enzymes, and Vb plus ATP is above all three under the present conditions. Because the enzymes were measured with different substrates, cofactors, oxygen tensions, and preparations, this comparison establishes catalytic capacity but not relative physiological flux.

A third comparison is with the other nitrite reductases of this field. For xanthine oxidase, the enzyme most often proposed as a mammalian nitrite reductase, Maia and Moura obtained a kcat of 0.693 s⁻¹ and a Km for nitrite of 0.585 mM, together with 0.244 s⁻¹ for the homologous aldehyde oxidoreductase of *Desulfovibrio gigas* [53]; Li et al. independently obtained an apparent Km for nitrite of 2.4 ± 0.2 mM for the same enzyme [54]. For deoxyhemoglobin, the reduction of nitrite to NO and methemoglobin proceeds with a rate constant of 2.9 M⁻¹·s⁻¹ at 25 °C and pH 7.0, and in deoxygenated whole blood exposed to 200 µM nitrite at 37 °C iron-nitrosylated hemoglobin formed with an observed rate constant of 0.0035 s⁻¹ [55]. The comparison with Table III can only be semiquantitative: the xanthine oxidase constant is a kcat obtained at saturation, whereas the values reported here were obtained at a single nitrite concentration and are lower bounds; and the hemoglobin reaction is second order in nitrite and heme and consumes ferrous heme stoichiometrically, whereas Cco is re-reduced through cytochrome c and turns over repeatedly. Within those limits the turnover rates in Table III are of the same order as the xanthine oxidase kcat, and for the Vb isozyme above it.

A second scale is provided by the oxidase capacity in Figure 4. The baseline rate of 0.95 µmol O₂·mg⁻¹·min⁻¹ equals 950 nmol O₂·mg⁻¹·min⁻¹. The Table III Cco/NO rates span 1.177–9.959 nmol NO·mg⁻¹·min⁻¹ (lowest, Va with ATP; highest, Vb with ATP) and therefore correspond to 0.12–1.05 NO molecules per 100 O₂ molecules of oxidase capacity. On an electron-equivalent basis, with four electrons consumed per O₂ and one per nitrite reduced to NO, this is approximately 0.031–0.26% of the oxidase electron capacity. These are capacities from different assay regimes, not a simultaneous electron partition: Table III was obtained in anoxia, whereas oxidase activity requires O₂.

The useful mechanistic distinction is therefore not enzymatic versus catalytic, because all enzymes are catalysts. Cco/NO is a respiratory-chain-coupled, multi-turnover nitrite-reductase activity that can be continuously regenerated by electrons delivered through cytochrome c. NOS enzymes are continuous NADPH-dependent catalysts, and Godber et al. showed that xanthine oxidoreductase also catalytically reduces nitrite using NADH or xanthine [56]. In contrast, the deoxyglobin reaction with nitrite consumes ferrous heme and forms ferric heme, so each turnover requires enzymatic re-reduction of the heme. Huang et al. demonstrated the allosterically controlled nitrite-reductase behavior of hemoglobin [57]. Electrons reaching Cco in intact cells also originate ultimately from NADH/FADH₂; the present assay bypasses most of the respiratory chain with ascorbate/TMPD and added cytochrome c.

The fractional contribution of Cco to total nitrite-derived NO is therefore expected to be context-dependent. Quesnelle et al. directly compared myoglobin and isolated mitochondria and found myoglobin to be the more efficient nitrite reductase at similar approximate concentrations [58]. In CHO cells lacking myoglobin, respiratory-chain inhibitors suppressed most of the nitrite-dependent signal, whereas expression of myoglobin increased NO formation and removed the significant inhibitor effect [58]. These findings support a potentially substantial mitochondrial contribution in cells or microdomains with little globin, but they argue against a universal claim that Cco dominates tissue NO production. The present study quantifies intrinsic turnover and capacity per amount of preparation; it does not determine a percentage of total NO in intact cells.

### 3.5. Both Cco/NO activity and ADP/ATP levels increase transiently when yeast cells experience a hypoxic shift

The above findings demonstrate that adenine nucleotides can modulate Cco/NO activity but they do not provide a physiological context in which changes in ADP or ATP levels may be relevant. To examine this, we have chosen to look at the effects of exposure to hypoxia on ADP and ATP levels and Cco/NO activity. We have observed previously that yeast cells experience a transient increase in protein tyrosine nitration during the first 4 hours after a hypoxic shift from normoxia to hypoxia then anoxia and speculated that this resulted from elevated Cco/NO activity [14]. Here, we have asked if Cco/NO activity *per se* increases during the first four hours of a hypoxic shift. For these studies, we used JM43, a wild-type yeast strain that contains a functional copy of the *YHB1* gene as well as functional copies of both *COX5a* and *COX5b*. Yeast cells were grown to mid-exponential phase in a fermentor sparged with air. They were then shifted to anoxia by sparging with O₂-free nitrogen gas containing 2.5% CO₂, as previously described [37], and samples were taken at different intervals after the shift and processed for measurement of ADP and ATP levels and of Cco/NO activity. The data in Figure 6A indicate that Cco/NO activity increases and peaks 120 min after the shift and then declines to its normoxic levels by 240 min. Over this 4 hour period neither the overall level of Cco nor mitochondrial respiration changes by more than a modest amount [59]. Measured independently, from respiration capacity and cytochrome content in whole cells of the same strain, the in vivo turnover rate of Cco rises transiently after a shift to anoxia, reaching a maximum about 20% above the normoxic value [59] within the first two hours — the same interval in which Cco/NO capacity peaks here — before declining at later times. The increase in Cco/NO capacity over that interval is considerably larger than 20%, so it cannot be accounted for by the change in oxidase turnover alone [59], suggesting that any increase in NO production by Cco/NO results from activation of Cco/NO. The increase in Cco/NO activity cannot be explained by replacing the aerobic subunit V isoform, Va, with the more active subunit V isoform, Vb, because subunit Vb does not appear in mitochondria until 8 hours after the shift [59]. To determine whether adenine nucleotides could account for the changes in Cco/NO activity during a shift, we measured cellular ADP and ATP levels. Figure 6B shows that the cellular ADP/ATP ratio rises transiently during a hypoxic shift, and that both of its components move: ADP increases and ATP decreases, each reaching its extreme value near 120 min and returning towards pre-shift values by 240 min. Because both terms change, the shift in the ratio cannot be attributed to a fall in ATP alone. The fall in ATP agrees with in vivo 3¹P-NMR measurements in *S. cerevisiae*, in which withdrawal of the oxygen supply lowered the intracellular ATP pool from 6.1 to 3.1 µmol (g dry weight)⁻¹ and the [ATP]:[ADP] ratio from 4.1 to 2.6 within 15 min [60], and with direct determinations in respiratory-competent cells, in which the cellular ATP/ADP ratio fell from 4.3 ± 0.2 with glucose in aerobiosis to 2.9 ± 0.3 with glucose in anaerobiosis [61]. The chemostat determination averaged spectra acquired 12.5 and 15 min after the shift [60], and the comparison in respiratory-competent cells was made between established steady states [61]; neither interval covers the excursion reported here, which develops over the first two hours. In the chemostat study the ADP level itself remained quasi-constant, so that the fall in the ratio was carried by the decrease in ATP [60]. In the measurement reported here both terms move, and the transition is sampled throughout the first 8 h in batch culture at mid-exponential phase rather than in a single window 15 min after the shift in a glucose-limited chemostat. Published values for the aerobic [ATP]:[ADP] ratio in this organism span two orders of magnitude, from 0.09 to 7.8 [60]. The rise in the diphosphate pool is documented as well: 3¹P-NMR of perchloric acid extracts of glycolysing *S. cerevisiae* showed the nucleoside diphosphate level to be about threefold higher anaerobically than aerobically in derepressed cells, though not in glucose-repressed cells [41]. That determination reports the total nucleoside diphosphate pool, a broader quantity than the ADP determination used here; the approximately threefold rise we measure nonetheless matches it. That the perturbation is confined to the transition rather than to the anaerobic end state is supported by the finding that the adenylate energy charge of *S. cerevisiae* is indistinguishable in anaerobic and fully aerobic steady states (0.83 ± 0.01) yet falls to 0.54 ± 0.03 at an intermediate oxygen supply of 2.8% [62]. AMP was not determined in these experiments, so the adenylate pool balance is not closed; adenylate not recovered as ATP or ADP would be expected to appear as AMP through the adenylate kinase reaction. Comparable changes in adenine nucleotide levels have been described in mammalian tissue and in plants exposed to low oxygen [63–66]. Interestingly, the increase in Cco/NO activity parallels an increase in the ratio of ADP to ATP during the shift (Figure 6B), consistent with a mechanistic link in which changes in intracellular ADP/ATP levels during exposure to hypoxia contribute to the changes in Cco/NO activity, although the parallel time courses establish association rather than proof of causation in vivo.

Previously, we have proposed that Cco/NO-generated NO and superoxide from mitochondrial complex III combine to form peroxynitrite, which then tyrosine nitrates protein components of a hypoxic signaling pathway [14, 45, 67]. The differential effects of ATP and ADP on Cco/NO and Cco/H₂O activity, together with the fluctuation in the ATP/ADP ratio during a hypoxic shift, suggest the following model (Figure 7) by which Cco transiently modulates both the generation of superoxide and NO in cells experiencing hypoxia. Insofar as high ATP/ADP ratios have been reported to depress the inner mitochondrial membrane potential and, by extension, the generation of free radicals [68, 69], the transient decrease in the ATP/ADP ratio during a hypoxic shift would function to increase superoxide production and the subsequent increases in yeast protein carbonylation and oxidative stress reported earlier [70]. Conversely, the transient decrease in ATP/ADP ratio during a hypoxic shift facilitates a transient increase in Cco/NO activity and NO production, making more NO available to combine with the increased superoxide and produce elevated levels of peroxynitrite, which is capable of both protein tyrosine nitration and protein carbonylation. Interestingly, the time frame for the increase in the ADP/ATP ratio and the increase in Cco/NO activity correlates well with the increases in protein tyrosine nitration and protein carbonylation [14, 70].

**Figure 7.**
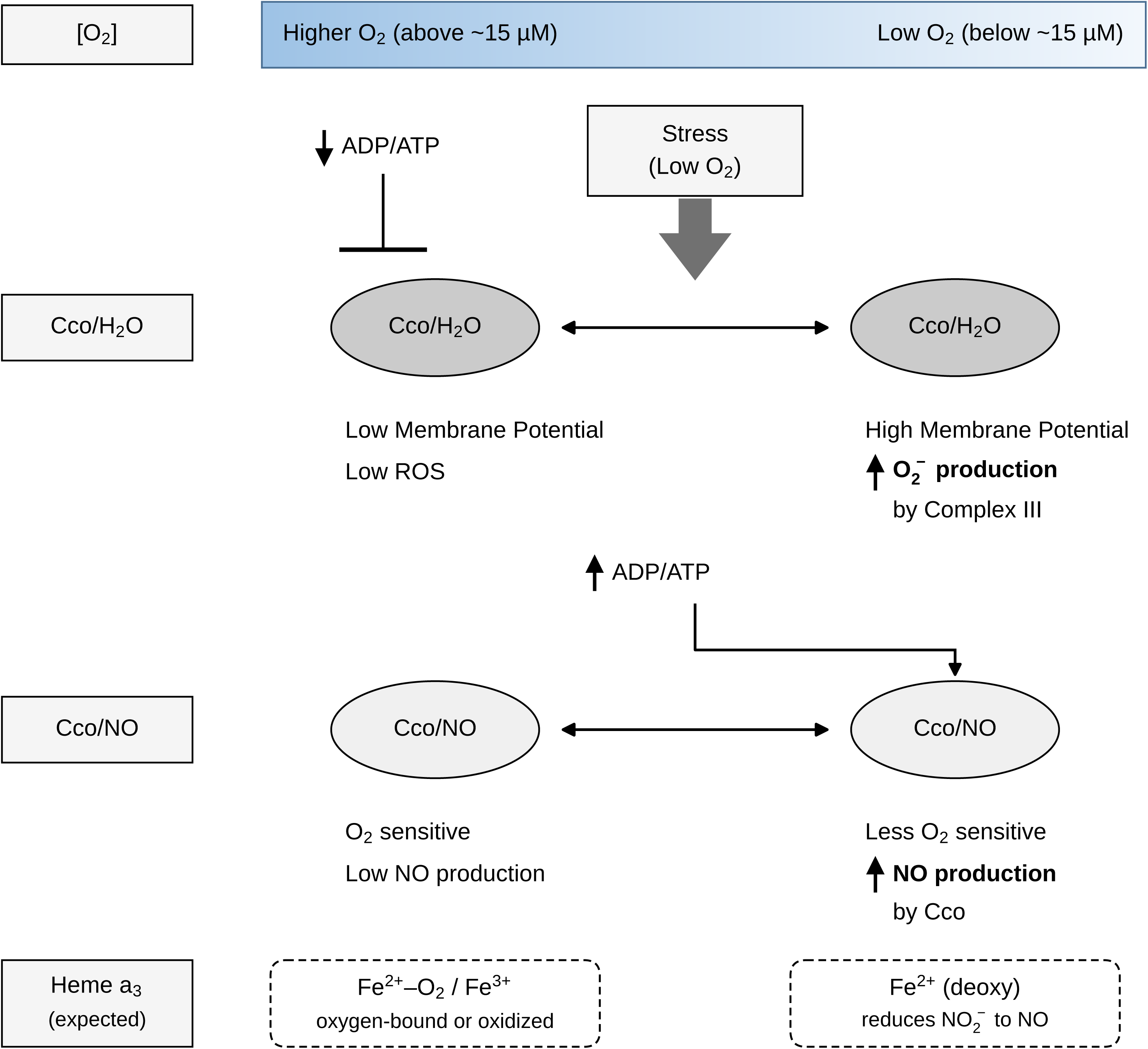
Model for coordinated regulation of Cco/H₂O, reactive oxygen species, and Cco/NO during hypoxic stress. The scheme summarizes the results in Figures 1–6 and is a model, not an additional experimental result. At a low ADP/ATP ratio, ATP-dependent inhibition of Cco/H₂O is associated with low membrane potential and low ROS, while Cco/NO remains oxygen-sensitive and NO production is low. During low-O₂ stress, an increased ADP/ATP ratio relieves inhibition of Cco/H₂O, increasing membrane potential and complex III superoxide production, and reduces the apparent oxygen inhibition of Cco/NO and increases NO production. In JM43 cells this shift is transient, with Cco/NO capacity and the cellular ADP/ATP ratio peaking near 120 min after the shift and returning towards pre-shift values by 240 min (Figure 6). To relate the scheme to the measured oxygen dependence, the low-ADP/ATP state corresponds to higher O₂ (above ∼15 µM), where the catalytic heme a₃ is expected to be predominantly in oxygen-bound or oxidized states (FeZ⁺–O₂ / Fe3⁺) and Cco/NO is suppressed, whereas the high-ADP/ATP state corresponds to low O₂ (below ∼15 µM), where a larger fraction of heme a₃ is expected to be in the reduced, deoxy-ferrous (FeZ⁺) state that supports nitrite reduction to NO. In the scheme, the shaded band indicates the dissolved O₂ condition and the dashed boxes give the expected heme a₃ iron state for each condition; these iron states are inferred from known heme chemistry and were not measured.

### 3.6. Cco/NO in cellular signaling

The findings presented here show that Cco/NO capacity is regulated across a wide range of oxygen concentrations and is strongly influenced by adenine nucleotides and subunit V isoform. Previous studies implicated Cco/NO in hypoxic signaling [14–16, 45, 67]. The present results provide a biochemical mechanism by which the increase in ADP/ATP accompanying hypoxia could enhance this activity. The data also show that ADP permits measurable Cco/NO activity at oxygen concentrations found in many mammalian tissues, but this observation was made with 1 mM nitrite in a detergent-solubilized preparation. It therefore establishes normoxic biochemical capacity rather than demonstrating Cco-derived NO production in normoxic intact cells. A cellular context in which this regulation can be tested already exists: yeast mutants that mimic caloric restriction show both an increased respiratory chain and increased NO levels, and their extended lifespan is NO-dependent [71]. That system couples the same two variables examined here, respiratory capacity and NO, in intact cells.

Assessing the biological plausibility of Cco/NO requires distinguishing enzyme-normalized rates from local NO concentrations. Mitochondria can metabolize nitrite and can also be targets of nitrite-derived NO [1, 72]. The rate measured here in mouse brain mitochondria at 30 µM O₂ with ADP (0.191 nmol NO·mg mitochondrial protein⁻¹·min⁻¹) cannot be compared directly with rates reported per 10⁶ human leukocytes [73] because the normalization bases differ. Nanomolar NO reversibly inhibits Cco in competition with O₂ [48], whereas estimated physiological NO concentrations near neuronal or endothelial sources fall within the sensitivity range of soluble guanylyl cyclase [74, 75]. Because NO reacts rapidly with superoxide [76], local NO and ROS concentrations near the inner mitochondrial membrane will help determine whether Cco/NO-derived NO acts primarily in signaling or contributes to peroxynitrite formation.

The isoform-specific ATP response is a distinctive result of this study. ATP reduces the Va turnover to 0.8 of Va baseline but increases Vb turnover 2.7-fold relative to Vb baseline; Vb plus ATP is 16.4-fold above Va baseline (Table III). A plausible hypothesis is that the molecular environment of a nucleotide-sensitive region differs between Va and Vb [28, 29, 77]. The cryo-EM structure of the yeast III₂IV₂ supercomplex provides structural context for Cox5A and its association with allosteric ATP inhibition [78], but it does not establish an equivalent Cox5B structure or mechanism. Direct comparison of the two isoforms, binding measurements, and site-directed mutagenesis will be required to test this hypothesis [30].

### 3.7. Experimental scope and unresolved physiological questions

All assays used 1 mM exogenous nitrite, two to three orders of magnitude above physiological tissue nitrite. Early measurements gave 12 ± 5 µM in rat heart [9]; later chemiluminescence determinations give 0.45–1.68 µM across brain, heart, liver, kidney and lung, and 22.5 µM in aorta [33]. The two sets of determinations differ by about an order of magnitude, and both are given here rather than the more favorable one; brain, the tissue used in the mammalian experiments reported here, is one of the two exceptions in which steady-state nitrite is highest [33]. On either estimate the concentration used here is far above the physiological range, and this is the principal limitation on extrapolating these rates to intact tissue. It is nonetheless worth recording how the authors of the lower estimate read their own number: Zweier et al. described it as a large pool of nitrite present in these hearts at the onset of ischemia, which could be reduced to form NO, and loading hearts with 10 µM nitrite increased post-ischemic injury and abolished most of the protection conferred by L-NAME [9]. Assaying a nitrite reductase well above the steady-state tissue concentration is also the standard practice of this field, both for signal-to-noise and because the enzymes of this class have millimolar Km values for nitrite [18]. For xanthine oxidase, the most extensively characterized of them, the Km for nitrite is 0.585 mM [53], 2.4 ± 0.2 mM in an independent determination [54], and 15.7 and 35.7 mM with xanthine and NADH respectively as reducing substrate [56]; Godber et al. noted explicitly that these constants are considerably higher than the micromolar nitrite concentrations found in plasma [56]. The 1 mM used here is of the same order as the lowest of those values and below the others. This constraint is organism-dependent: mammals lack an effective nitrate reductase and depend on the entero-salivary circulation for nitrite supply [13], whereas plants reduce nitrate to nitrite endogenously and accumulate nitrite under oxygen deficiency, conditions under which mitochondrial nitrite reduction to NO is well documented [79, 80]. These substrate-driven experiments define Cco capacity under controlled conditions. For yeast the relation between the concentration supplied and the one the enzyme encounters has been measured, although it applies to intact preparations rather than to the solubilized one used here. In intact mitochondria only about 10% of externally supplied nitrite is internalized, and the intracellular nitrite concentration of aerobic JM43 cells was determined to be 9.5–17.5 nmol (g wet weight)⁻¹, corresponding to 15–27.5 µM; in the same work NO formation was detectable with as little as 20 µM internalized nitrite, a value within that measured cellular range [14]. That the reaction proceeds at physiological nitrite is therefore established, but by those earlier intact-mitochondria experiments and not by the present ones. In the solubilized preparation used here there is no membrane barrier, and the enzyme is exposed to the full 1 mM. Nitrite was not titrated in this preparation, and 1 mM is not established to be a saturating concentration for Cco: with purified yeast Cco at pH 6.0 the rate rose from 1.3 to 14 to 221 nM NO·min⁻¹ as nitrite was raised from 0.02 to 0.1 to 1 mM, with no approach to a plateau [14], and a comparable concentration dependence was obtained in solubilized mitochondria [16]. The rates reported here are therefore capacities measured at one fixed substrate concentration rather than physiological fluxes, and they are lower bounds on that capacity rather than maximal velocities. One consequence is worth noting. Across the fiftyfold range from the micromolar concentrations at which the reaction was first detected to the 1 mM used here, oxygen consumption is unaffected. That is the behavior of a substrate, not of an inhibitor. What remains undetermined is the apparent Km of Cco for nitrite, the local concentration within the mitochondrial matrix, and whether intracellular nitrite changes during a hypoxic shift. One regime was not examined at all. Every experiment reported here was performed with nitrite in large excess over the enzyme, whereas the converse condition, in which reduced Cco is in excess over the nitrite available to it, is the one implied by submicromolar tissue nitrite. Whether the reaction remains multi-turnover in that regime, or becomes limited by the supply of nitrite, is not addressed by these data. A further consequence of working with independent mitochondrial preparations is that the absolute specific activity varies between them, so that the nucleotide-free rate against which each experiment is normalized is not a constant of the system: it is 0.5 nmol NO·mg protein⁻¹·min⁻¹ in the preparation used for Figure 2A and 2B, and 1.472 nmol NO·mg protein⁻¹·min⁻¹ for the Va strain in Table III. Every percentage and every relative turnover reported here is therefore referred to the nucleotide-free control of its own experiment, and rates from different figures should not be compared in absolute terms. Likewise, the 5–20 mM externally added nucleotide concentrations should not be interpreted as measured free matrix concentrations.

The Cco/NO assay used detergent-solubilized mitochondria and an artificial electron-donor system throughout, for the reason given in Experimental Procedures: externally supplied cytochrome c cannot reach Cco in an organelle with an intact outer membrane. The same choice that makes the measurement possible therefore also fixes what it can report. Respiratory control ratio, proton pumping, membrane potential and pH dependence were not measured, so the results speak to catalytic capacity rather than to operation in an energetically coupled organelle. Figure 6 likewise reports capacity, assayed in mitochondrial preparations from cells sampled during a hypoxic shift, and its correlation with ADP/ATP is evidence of association rather than of cause. Neither point leaves the intact-cell question open. Nitrite-dependent NO production has been measured directly in intact anoxic yeast cells, with in vivo turnover rates calculated against intracellular cytochrome aa₃ content: rates were abolished in ρ⁰ strains lacking a functional respiratory chain, were higher in strains lacking YHB1, and were 4.5-to 7-fold higher for the Vb isozyme than for the Va isozyme — the same isoform ranking obtained here in solubilized mitochondria [16]. A fraction of the NO generated by yeast mitochondria is also released from the cell [67]. What has not been done, and is the natural next step, is an intact-cell NO time course across the hypoxic shift under controlled nucleotide conditions. That experiment is now technically accessible: intracellular NO can be quantified with a detection limit of 6 nM by staining cells with the NO-specific probe DAF-FM DA and resolving the resulting triazole by high-performance liquid chromatography with fluorescence detection [81]. The same work shows why it must be measured in that way rather than from fluorescence intensity alone, since in yeast exposed to ethanol, vanillin or heat shock the probe-dependent fluorescence proved not to derive from NO [81]. Two features of this system favor the experiment: *S. cerevisiae* carries no orthologue of the mammalian NO synthase [82], and the strains used here lack *YHB1* and therefore the dominant NO-consuming activity. The adenine nucleotide values in Figure 6B are whole-cell determinations rather than free matrix concentrations, and the interpretation advanced here rests on the direction and relative magnitude of the change rather than on the absolute ratio at its peak. Two consequences follow from the solubilized preparation itself. Acid-catalyzed disproportionation of nitrite yields NO without any enzyme, and hypoxia acidifies the cytosol; the controls in Figure 1 exclude that route here, since no NO forms with heat-denatured mitochondria or without reductant and formation is abolished by azide, but in intact cells hypoxic acidification together with the transmembrane proton gradient could favor it. Acidification also acts on the enzyme-catalyzed route, and two independent systems place its onset near the same pH. With purified yeast Cco at 1 mM nitrite the rate rose from 1.0 to 17 to 221 nM NO·min⁻¹ as the assay pH was lowered from 7.0 to 6.5 to 6.0 [14]; in ischemic rat heart, myocardial pH falls to 5.9 within 10 min and the threshold pH at which NO generation was first seen was approximately 6.0 [9]. The Assay Buffer used here is 200 mM Tris-Cl at pH 7.0, the unfavorable end of that dependence, which is a further reason to read the present rates as lower bounds. Conversely, because the preparation lacks a functional F₁F₀ ATP synthase, the inhibition of oxidase activity by ATP reported here is necessarily allosteric: it cannot arise from ATP hydrolysis or from changes in the proton-motive force. In intact mitochondria both mechanisms would operate together.

Cco both produces and reacts with NO at the heme a₃–Cu_B_ center [19–23]. Binding, oxidation, or re-consumption of newly formed NO could therefore make the electrode signal a net escape rate rather than the gross chemical formation rate. The same chemistry raises a question that the present results invite. If Cco reduces nitrite to NO, and NO inhibits Cco at nanomolar concentrations in competition with oxygen [48], why is the oxidase not chronically inhibited in tissues where nitrite is present at one to twenty micromolar? Four considerations bear on this. First, NO formation is concentration-dependent, and the rate at micromolar nitrite is far below the rate measured here at 1 mM [14], so the steady-state NO concentration it can support is correspondingly low. Second, NO is consumed by several routes at once: by the globins — YHb in yeast, myoglobin and hemoglobin in mammalian tissue — and by Cco itself, which rapidly metabolizes NO back to nitrite [21, 22]. The product of the reaction is therefore also its substrate, and the enzyme does not accumulate a stable inhibitory complex while it is turning over. Third, a mild and reversible restraint on oxygen consumption at low oxygen tension is not a malfunction but the proposed physiological role of the nitrite pool [13]. Fourth, and most directly, the activity reported here is transient: Cco/NO capacity peaks near 120 min after a hypoxic shift and returns to its pre-shift value by 240 min (Figure 6A). An activity that is switched on and off within hours can act as a signal; one that were constitutive could not. The effect of nitrite on oxidase activity across a full oxygen series was not systematically quantified. Deletion of YHB1 removes a major NO-consuming pathway and is useful for isolating formation at the Cco step, but it also prevents extrapolation to the complete NO balance of wild-type yeast.

Taken together, these data establish that Cco/NO is a metabolically and isoform-regulated, multi-turnover nitrite-reductase activity. Assigning it a fractional share of total cellular or tissue NO is a separate question, and will require matched measurements in intact cells or coupled mitochondria at physiological nitrite, with total NO flux compared against Cco-specific genetic or pharmacological perturbation and the abundance and activity of NOS, globins, xanthine oxidoreductase, and other competing sources and sinks.

### 3.8. Yeast as a model for the mammalian enzyme

These experiments were performed in *Saccharomyces cerevisiae* not because yeast is the intended endpoint but because the regulatory architecture under study is conserved. The oxygen-regulated subunit V isoform pair of yeast Cco has a direct mammalian counterpart in the COX4-1 and COX4-2 isoforms of subunit IV [77], and the yeast system is what led to its discovery. In demonstrating oxygen-regulated isoform switching in mammalian cells, Fukuda et al. begin from the observation that in yeast COX subunit composition is regulated by *COX5a* and *COX5b* transcription in response to high and low O₂ respectively, and then show that hypoxia-inducible factor 1 reciprocally regulates COX4 subunit expression by activating transcription of the genes encoding COX4-2 and LON, the mitochondrial protease required for COX4-1 degradation [83]. The subunit V/IV isoform switch is therefore not a fungal idiosyncrasy but an oxygen-sensing device that yeast made tractable and mammals subsequently confirmed. For the same reason, a nucleotide- and isoform-dependent gating of nitrite reduction established in the yeast enzyme constitutes a testable prediction about the mammalian enzyme rather than an observation confined to fungi. The mouse brain mitochondria used here approach the same question from the other direction, and the two preparations respond to ADP in the same way.

### 3.9. Physiological relevance

Five independent lines of evidence indicate that Cco/NO is a credible contributor to cellular NO rather than an in vitro curiosity. First, its enzyme-normalized turnover lies in the same range as that of the purified NOS isoforms — 0.43–8.74 NO molecules·Cco⁻¹·s⁻¹ here, against approximately 0.21–0.27 s⁻¹ for eNOS, 3.2 s⁻¹ for nNOS and 2.4–2.8 s⁻¹ for iNOS [50–52] — and the Vb isoform in the presence of ATP numerically exceeds all three under the nitrite-saturating conditions of these assays. Because the rate rises steeply with nitrite concentration — 170-fold, from 1.3 to 221 nM NO·min⁻¹, over a 50-fold rise from 0.02 to 1 mM with purified yeast Cco at pH 6.0 [14] — rates at physiological nitrite (1–20 µM) are expected to be lower than those reported here. The magnitude of that reduction cannot be established from the available measurements, so this comparison concerns catalytic capacity rather than physiological flux. Second, Cco sustains multi-turnover catalysis: it is continuously re-reduced through cytochrome c, and, having no stable ferrous–dioxygen species to scavenge the NO it forms and a comparatively low affinity for NO even at its ferrous heme, it is not trapped as a nitrosyl complex during turnover but instead releases nitrite and cycles again [21, 22]. The globins are themselves re-reduced enzymatically, by cytochrome b5 and cytochrome b5 reductase [84], so continuous reduction is not what distinguishes the two systems; the distinction lies in the stability of the ferrous–dioxygen state and in the affinity for the NO produced. Third, in intact cells lacking myoglobin the mitochondrial contribution is dominant: inhibitors of the respiratory chain abolished approximately 80% of nitrite-dependent NO formation in CHO cells, and this inhibitor effect disappeared when myoglobin was expressed [58]. Because myoglobin is restricted to cardiac and skeletal muscle, most tissues — including brain, liver, kidney and vascular endothelium — correspond to the myoglobin-free condition, and nitrite-dependent mitochondrial S-nitrosation has been reported in liver and brain [58]. That modification is distinct from the tyrosine nitration discussed above: S-nitrosation adds a nitroso group to a cysteine thiol and is reversible, whereas nitration adds a nitro group to a tyrosine ring and is not, so the two report different fates of the same NO. Fourth, the activity has been demonstrated in intact anoxic yeast cells and not only in isolated or solubilized preparations, with rates abolished in respiratory-chain-deficient ρ⁰ strains and with the same Va/Vb isoform ranking obtained in vitro [16], and a fraction of the NO produced is released from the cell [67]. Fifth, the activity is gated by adenine nucleotides and by subunit V isoform, and ADP shifts the apparent oxygen sensitivity of Cco/NO toward lower O₂ values, with statistically significant effects at 10 µM O₂ and below; a reaction under this degree of metabolic control is more readily interpreted as a regulated signaling output than as an incidental side reaction. Consistent with this, yeast mutants with an increased respiratory chain also show increased NO levels and an NO-dependent extension of lifespan [71]. Together with the conservation of respiratory-chain-dependent nitrite reduction across yeast, plants, algae and mammals [45], these considerations place Cco/NO among the candidate nitrite reductases in tissues where globins are scarce, and identify the ADP/ATP ratio as one determinant of its catalytic output.

Finally, because most experiments reported here used detergent-solubilized mitochondria, which lack inner-membrane integrity and therefore have neither a proton gradient nor a membrane potential, the effects of the ATP/ADP ratio measured here are properties of the enzyme and do not necessarily reflect the net effect of the cellular ADP/ATP ratio in intact, energetically coupled mitochondria in vivo.

## ACKNOWLEDGEMENTS

We thank Pamela S. David for the purification of yeast cytochrome c oxidase, Dong Kyun Woo for the construction of the yeast strains YDW27 and YDW29, and Gretchen H. Stein for providing the mouse forebrain tissue used in this work. We also thank Michael Stowell for the gift of mice and Margaret Isenhart for her assistance with the animal facility.

## CREDIT AUTHORSHIP CONTRIBUTION STATEMENT

P.R.C.: Conceptualization, Methodology, Investigation, Validation, Formal analysis, Data curation, Visualization, Writing – original draft, Writing – review & editing. K.A.B.: Methodology, Investigation, Validation, Writing – review & editing. R.O.P.: Conceptualization, Funding acquisition, Supervision, Writing – original draft, Writing – review & editing.

## DATA AVAILABILITY

All data generated or analysed during this study are included in this article.

## DECLARATION OF GENERATIVE AI AND AI-ASSISTED TECHNOLOGIES IN THE MANUSCRIPT PREPARATION PROCESS

During the preparation of this work the authors used Claude (Claude Opus 5, Anthropic) to assist with language editing, with the organization and drafting of portions of the text, with verification of the cited literature against the primary sources, and with checking the internal numerical consistency of the tables. No experimental data were generated, analyzed or altered by these tools, and no figures were produced or modified by them. After using this tool the authors reviewed and edited the content as needed and take full responsibility for the content of the publication.

---

**Supplementary Table S1.** Individual recordings of net NO production by mouse brain mitochondria at each O₂ concentration (nmol NO·mg protein⁻¹·min⁻¹)

|  | + ADP |  |  |  | + ATP |  |  |  | No nucleotide |  |
|---|---|---|---|---|---|---|---|---|---|---|
| O<sub>2</sub> (μM) | 1 | 2 | 3 | 4 | 1 | 2 | 3 | 4 | 1 | 2 |
| 175 | 0.0259 | 0.0439 | 0.0476 | 0.0442 | 0.0357 | 0.0332 | — | — | — | 0.0061 |
| 150 | 0.0450 | 0.0586 | 0.0503 | 0.0451 | 0.0318 | 0.0242 | — | — | — | 0.0029 |
| 100 | 0.0728 | 0.0773 | 0.0543 | 0.0577 | 0.0137 | 0.0049 | 0.0122 | 0.0379 | 0.0265 | −0.0218 |
| 75 | 0.0969 | 0.0921 | 0.0669 | 0.0709 | 0.0055 | −0.0029 | 0.0061 | 0.0318 | 0.0233 | −0.0303 |
| 50 | 0.1441 | 0.1202 | 0.1016 | 0.1138 | −0.0043 | −0.0066 | 0.0026 | 0.0265 | 0.0201 | −0.0383 |
| 40 | 0.1753 | 0.1367 | 0.1278 | 0.1498 | −0.0049 | −0.0088 | 0.0024 | 0.0253 | 0.0191 | −0.0439 |
| 30 | 0.2215 | 0.1642 | 0.1720 | 0.2069 | −0.0083 | −0.0135 | 0.0012 | 0.0229 | 0.0180 | −0.0448 |
| 20 | 0.3046 | 0.2121 | 0.2514 | 0.3047 | −0.0100 | −0.0159 | −0.0018 | 0.0218 | 0.0167 | −0.0497 |
| 15 | 0.3872 | 0.2509 | 0.3170 | 0.3772 | −0.0113 | −0.0147 | −0.0022 | 0.0195 | 0.0216 | −0.0490 |
| 10 | 0.5442 | 0.3105 | 0.4324 | 0.4862 | −0.0158 | −0.0184 | −0.0035 | 0.0190 | 0.0236 | −0.0523 |
| 5 | 0.9330 | 0.4227 | 0.6592 | 0.6601 | −0.0054 | −0.0184 | 0.0044 | 0.0260 | 0.0446 | −0.0521 |
| 0 | 1.0249 | 1.0555 | 1.1887 | 1.5664 | 0.2098 | 0.2177 | 0.2731 | 0.2473 | 0.3225 | 0.2856 |

Values as recorded in the original data files; each column is one recording. With ADP, recordings 1–2 and 3–4 were made on two different days; likewise with ATP. Rates are slopes of the NO-electrode trace and are therefore net rates (formation minus consumption); negative values correspond to a falling NO signal, from NO consumption or electrode baseline drift. —, O<sub>2</sub> concentration not covered by that recording. Means ± standard error and the criterion for n.d. are given in Table II.

